# A natural temperature, elevation, and rainfall gradient over the Cascade Mountains drives changes in *Populus trichocarpa* stress response and microbial community structure

**DOI:** 10.64898/2025.12.04.692393

**Authors:** Kathryn E. Bazany, William Argiroff, Alyssa A. Carrell, Kelsey R. Carter, Nancy L. Engle, Sara Jawdy, Dawn M. Klingeman, Justus Smith, Ian Morris, Amy Schaefer, John Lagergren, Daniel Jacobson, Stanton L. Martin, Melissa A. Cregger, David J. Weston, Dale A. Pelletier, Christopher W. Schadt

## Abstract

- Abiotic stresses associated with warming and drought are increasing in frequency and severity due to climate change. However, microbial associations can partially mitigate their adverse effects on plant physiology.
- We examined the bacterial/archaeal and fungal communities with 16S rRNA and ITS2 gene region amplicon sequencing of bulk soil, rhizosphere soil, and root endosphere samples of *Populus trichocarpa* trees from 11 sites along a 200 km transect with highly variable precipitation, temperature, and soil conditions over the Cascade Mountains in Washington, USA. Foliar stress response was examined with qRT-PCR and field physiological measurements. Soil conditions were examined with physiochemical sample analysis and site climate conditions were approximated using DAYMET.
- For each niche sampled, bacterial/archaeal and fungal communities clustered into three distinct micro-climate groups: a high-precipitation, low-stress community on the western side of the range, and two stressed climate communities––a high-elevation community nearer the crest of the pass, and a warmer and drier community along the eastern slope.
- The differences between these micro-climate communities, though still distinct, were notably diminished within the *P. trichocarpa* root endosphere, indicating that *P. trichocarpa* likely has a homogenizing effect on the microbiome under stress, but climate plays a major role in shaping these microbial communities.

## Introduction

Increasing plant abiotic stressors like drought and rising temperatures can drive broad ecosystem changes (Hartmann *et al*., 2022), including altering the microbial communities of soil and plants (Baldrian, López-Mondéjar and Kohout, 2023). Climate and soil physiochemical properties influence the microbial communities of soil, while microbial diversity in turn influences soil functioning, which has broad implications for ecosystems and climate feedback loops (Lange *et al*., 2015; Crowther *et al*., 2019; Delgado-Baquerizo *et al*., 2020; Bastida *et al*., 2021; Spohn *et al*., 2023). Temperature, moisture, pH, soil structure and nutrient status all influence soil microbial community composition by setting the limits of microbial survival and driving competitive advantages for certain taxa (Delgado-Baquerizo *et al*., 2018; Wu *et al*., 2022; Knight *et al*., 2024). Plant-associated microbial communities are also affected by abiotic conditions, both directly, as the soil microbial community is the primary source of microbes that colonize the plant environment (Dove *et al*., 2021) and indirectly, as plants tailor their microbiomes to better respond to stress (Trivedi *et al*., 2020). Plants selectively enrich certain microbes in their rhizosphere, the region of soil under the influence of the root, via the release of exudate carbon and other signaling molecules (Broeckling *et al*., 2008; Chaparro *et al*., 2013; Schaefer *et al*., 2013; Coutinho *et al*., 2018; Hu *et al*., 2018). Select microbes from the rhizosphere enter and colonize the root compartment by passing physical and plant immune system barriers (Hacquard *et al*., 2017; Teixeira *et al*., 2021). The plant endosphere is also subject to priority effects, which introduce additional stochasticity to microbial community assembly as determined by order of arrival (Dove, Veach, *et al*., 2021; Debray *et al*., 2022). As a result, microbial community environments may respond directly and indirectly to abiotic stressors, and this response may vary greatly under different habitat and host factors.

Due to the complexity of soil and plant-associated microbial communities, predicting their responses to changing environmental conditions, as well as making meaningful predictions as to how plant and microbial community changes will in turn influence the climate and ecosystem resilience, is an ongoing challenge. Experimental systems where warming, osmotic stress, and increased CO_2_ treatments are applied offer effective methods to address these questions, however they are expensive to establish, and it is impractical to determine how these plant-microbial systems will respond and adapt over the extended timeframes relevant to long-lived forest tree species. Naturally occurring climate gradients present a unique opportunity to examine the extended effects of climate variability on soil, plants, and plant-microbiomes. Gradient research may elucidate how microbiomes will respond to climatic shifts and help alleviate plant stress.

*Populus trichocarpa* is an ecologically important early-succession tree species with a broad geographic range (Cooke and Rood, 2007). It primarily grows in vital riparian habitats along streams and rivers and is tolerant to both flooding and drought due to its deep root networks. It is also economically important for biofuels and paper and fiber production due to its rapid growth rate (Tuskan *et al*., 2006; Gudynaitė-Franckevičienė and Pliūra, 2021). The *P. trichocarpa* genome has been fully sequenced (Tuskan *et al*., 2006) and extensive genome wide association study (GWAS) resources have been developed to identify genes linked to specific traits like abiotic stress response pathways and pathways for targeted microbial enrichment (O’Banion *et al*., 2023). This makes *P. trichocarpa* an ideal ecological model species to understand plant-microbiome interactions in trees (Cregger *et al*., 2021).

In this study, we established 11 sites along the Cowlitz and Tieton River watersheds over the Cascade mountain range in Washington State, USA. Rain shadow effects create a precipitation gradient where the Cowlitz watershed on the western slopes of the Cascades has significantly higher annual rainfall than the more arid Tieton watershed on the eastern side. Temperature also varies considerably, as high-elevation sites are cooler than the low-elevation sites on either side of the range. Soil conditions are however somewhat variable, as nutrients such as calcium, sodium, and iron leach at higher altitudes and rates of precipitation, and there are often reduced decomposition rates at high-elevation that results in higher carbon-to-nitrogen ratios.

We hypothesized that (1) the bacterial/archaeal and fungal communities would vary with the given microbial niches sampled (bulk soil, rhizosphere soil, and root endosphere) and among sites based on their variable environments, (2) *P. trichocarpa* would exhibit greater signs of abiotic stress at high and dry sites compared to wet sites, and (3) within niche habitats that specific climate, soil, and host factors would correlate with microbial community diversity, composition, patterns of co-occurrence, and assembly processes.

To test these hypotheses, we collected bulk soil, rhizosphere soil, and root endosphere samples from three *P. trichocarpa* trees at each site, then examined the bacterial/archaeal and fungal communities of these samples with 16S and ITS2 amplicon sequencing respectively. We also evaluated the soil physiochemical properties, leaf ecophysiological status, and the foliar stress response of each tree via a panel of abiotic stress related genes, as well as the climate status of each site.

## Methods

### Field Site Selection and characterization

Potential sampling sites were chosen based on environmental stress gradients computed with air temperature and soil water content following (Combs-Giroir *et al*., 2025). After field ground truthing in late June 2023, final sites were based on broad soil characteristics, the number/diameter of sampleable *P. trichocarpa* trees in 10m radius plots, and other practical considerations (S1). 11 total sites were sampled over the White Pass along the Cowlitz and Tieton River basins in Washington spanning roughly 200km in distance and 1300m in elevation (Figure 1). An additional transect of sites over McKenzie Pass in Oregon was also planned for sampling, however wildfires prevented us from proceeding with sampling.

**Figure 1:**
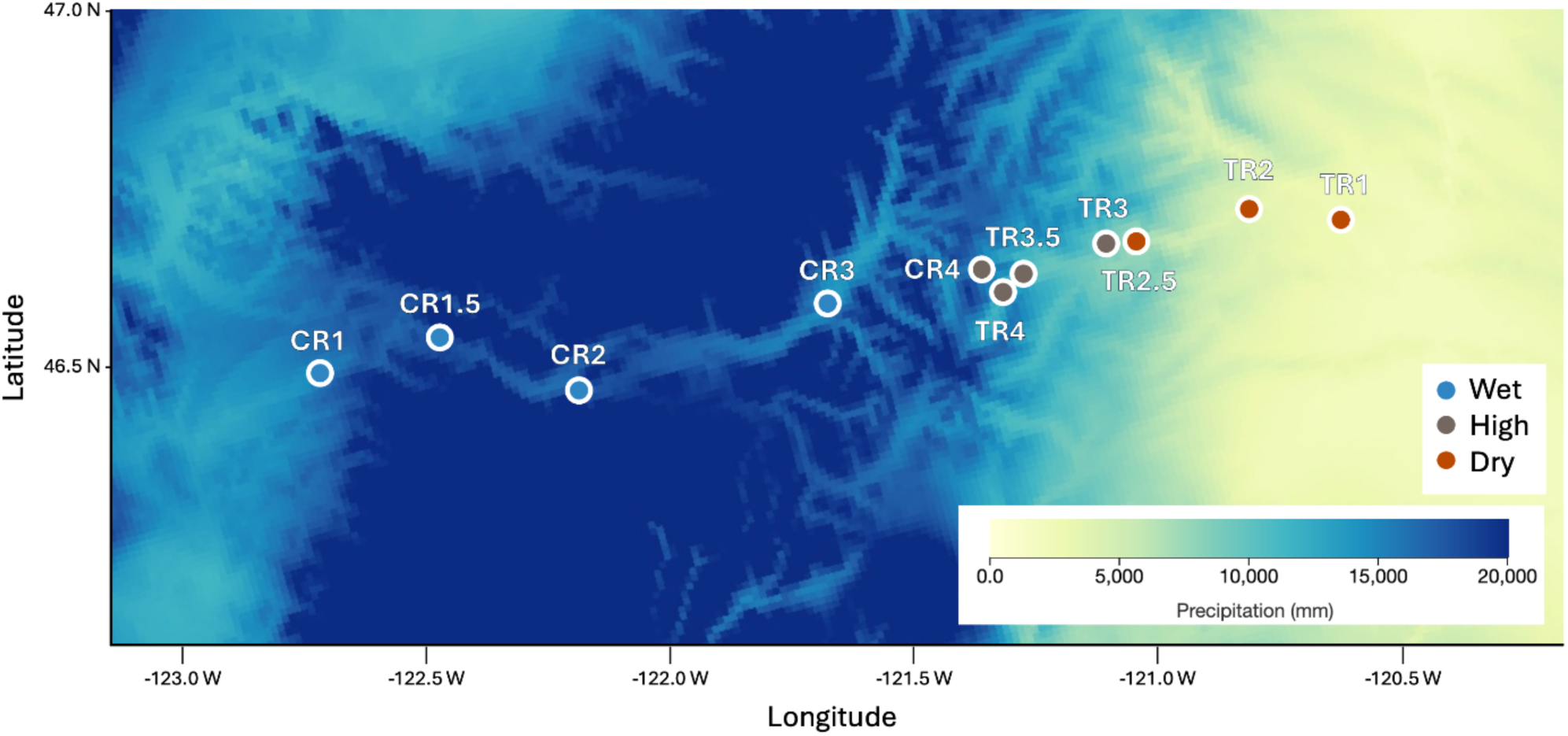
Site locations and total precipitation (mm) in the decade leading up to sampling (September 6, 2013 – September 6 – 2023) from DAYMET at a resolution of 1 km^2^. Sites are color coded based on beta-diversity clustering patterns and labeled by broad climate characteristics, blue for the high precipitation, wet sites, grey for the cool, high elevation sites, and orange for the low precipitation, dry sites.

Precipitation and temperature data were collected for each site from DAYMET (Thornton *et al*., 2022), a system of algorithms that interpolates and extrapolates climate data at an approximately 1 km^2^ resolution across the United States (Figure 1, S2). We collected and averaged precipitation and temperature data for each site from three time-frames, one month, one year, and one decade leading up to the sampling date. As all timeframes were highly correlated, we used the decadal averages for all models. Average maximum and minimum temperatures in all timeframes sampled were inversely correlated with site elevation (S2-3).

### Sample Collection and Tree Ecophysiological Characterization

Two soil cores at 5-15 and 40-50 cm depths were taken from each site for a total of four samples which were analyzed for total organic matter, carbon, nitrogen, phosphorus, iron, calcium, copper, potassium, magnesium, manganese, and sodium, as well as pH, cation exchange capacity, and soil, silt, and sand composition using standard soil analysis methods at the University of Georgia (UGA) Agricultural and Environmental Soil Analysis Lab. At CR1.5 only two soil samples, both at 5-15 cm depth were collected for soil chemistry analysis due to rocky conditions and was excluded from the soil physiochemical analyses. Soil status varied along the transect with some variables correlating with elevation, like pH, and C:N ratios, and others varying with watershed, like sand, silt, and clay content (S2-3).

Three *P. trichocarpa* trees were selected to be as close to the average diameter of all the trees in each plot and were sampled in early September 2023 near the end of the seasonal drought period in that region. To categorize the soil and root bacterial/archaeal and fungal communities, we collected a bulk soil, rhizosphere soil, and root samples. Bulk soil cores were taken from 0 to 20 cm depth near each tree sampled. Rhizosphere and root endosphere samples were taken from shallow roots excavated from each tree by tracing larger roots to branches containing fine roots which were collected along with the rhizosphere soil adhered to them. After the photosynthetic measurements, sampled leaves were collected for qRT-PCR analysis. Samples were transported to Oak Ridge National Laboratory in Tennessee on dry ice and stored at −80 C.

Plant physiological measurements were taken from canopy, sun exposed leaves. Two branches per tree were collected using a sling shot and immediately recut and placed in water. We measured gas exchange and fluorescence of three trees per plot, collecting 4-6 measurements between the two branches from each tree. Stomatal conductance (*g_sw_*), the photosynthetic efficiency of photosystem II (ΦPSII), leaf to air vapor pressure deficit (VPD_leaf_), and ambient light (Qamb) in micromoles of photons per square meter per second (µmol m⁻² s⁻¹) was measured using a handheld Porometer/Fluorometer (LI-600, LI-COR Inc., Lincoln NE, USA). Although we tried to minimize this by moving to an open area, some readings may have been influenced by shading or cloud cover.

### Sample Processing

DNA was extracted from the bulk soil and rhizosphere soils using the PowerSoil DNA isolation kit (Qiagen). Rhizosphere soils were defined as soil clinging tightly to the roots and were separated and processed according to protocols outlined by (Simmons *et al*., 2018) and modified by (Cregger *et al*. 2018). Fine roots (< 2 mm in diameter) were sterilized with protocols described in (Henning *et al*., 2016), frozen in liquid N, and ground into a fine powder using a bead-beater (Qiagen, Venlo, the Netherlands). DNA was isolated from the root samples using the PowerPlant Pro DNA isolation kits (Qiagen). DNA yield from each extraction was quantified on a NanoDrop 1000 Spectrophotometer (NanoDrop, Wilmington, DE, USA) and Qubit (ThermoFisher, Waltham, MA).

Archaeal/bacterial and fungal libraries were prepared following the Illumina 16S metagenomic sequencing library preparation guide (Part 15044223 Rev. B, Illumina, San Diego, CA). Primary amplification used custom 515F and 806R primers for 16S with PNA oligos added to the master mix to prevent mitochondria and chloroplast amplification and ITS2 gene region primers for fungi (Dove *et al.,* 2020; Rogers *et al.,* 2018), with subsequent barcoding using Illumina Nextera XT v2 indexes (S4). Paired-end sequencing (2×251×8×8) was performed on an Illumina MiSeq instrument (Illumina, San Diego, CA) using v2 chemistry.

RNA was extracted with the Promega Maxwell® RSC Plant RNA kit (Promega) with a few customizations (S5). RNA was synthesized into cDNA which was diluted 1:10 for qRT-PCR. Primers and cycling conditions for genes involved in abiotic stress response including genes involved in the ethylene, jasmonic acid, salicylic acid, and general stress and detoxification were tested and optimized on *Populus* foliar samples from a prior greenhouse drought experiment. Final qRT-PCRs were run on a BioRad Real-Time PCR System (S5). Relative expression was determined against 18S expression and UBQ10B expression as housekeeping standards and the average expression in the wet sites was used as a treatment control.

### Bioinformatics

Sequencing data was processed in R using DADA2 (Callahan *et al*., 2016). Sequencing primers were trimmed using CutAdapt (Martin, 2011), and filtered to a maximum of one base pair error. Forward and reverse reads were trimmed at 230bp and merged with a maximum mismatch of two base pairs. ASV tables were generated, and chimeric sequences were removed with DADA2 (Callahan *et al*., 2016). Taxonomies were assigned to the 16S reads by aligning to SILVA v.138.2 (Quast *et al*., 2013) and the ITS reads with UNITE v.10.0 (Abarenkov *et al*., 2024). 16S taxonomy assignments were filtered to remove chloroplast and mitochondrial reads. Reads that were unidentified at either the phylum or kingdom level only were also removed from both the 16S and ITS libraries.

### Data Analysis

All analyses were performed in R v.4.5.2. All code is available on github (https://github.com/kbazany/washington_transect_amplicon). For plant physiology analysis, linear mixed effects models (LMEs) were implemented to evaluate the strength of the correlations considering multiple samples within trees using lme4 (Bates, *et al*., 2015) and lmetest (Zeilies and Hothorn, 2002). Analysis of variance models (ANOVAs) were used in addition to the LMEs to evaluate the influence of site and climate type on the plant physiology metrics. For soil physiochemistry analysis, LMEs were used to examine the correlation between climate variables and soil factors and ANOVAs were used to examine the impact of site and climate category.

Shannon indexes were calculated with the vegan package v. 2.6-4 (Oksanen *et al*., 2025) to examine bacterial/archaeal and fungal alpha-diversity. LMEs were employed to assess the impact of continuous variables related to climate, soil, and host data. To assess the impact of categorical variables like niche and site on the bacterial/archaeal and fungal beta-diversity, we performed PERMANOVAs using the adonis2 function and non-metric multidimensional scaling (NMDS) ordination based on Bray-Curtis distances using the metaMDS function in vegan. In the NMDS ordination, sites clustered cleanly into three broad climate categories––wet sites (CR1, CR1.5, CR2, CR3), high altitude sites (CR4, TR4, TR3.5, TR3), and dry sites (TR2.5, TR2, TR1).

For continuous variables, including climate data, soil data, and tree phenotype data, we ran linear models in adonis2 and visualized the results with canonical correspondence analysis (CCA) plots in vegan (Oksanen *et al*. 2025). To evaluate patterns of enrichment and depletion among bacterial/archaeal and fungal phyla we performed site-wise analysis of compositions of microbiomes with bias-correction (ANCOM-BC) (Lin and Peddada, 2020).

Co-occurrence networks were generated using SpeicEasi (Kurtz, 2025) and iGraph (Csárdi *et al*., 2025) in R. Data was subset by niche into bulk soil, rhizosphere soil, and root endosphere, then within each niche into wet, high, and dry sites consistent with ordination results. This resulted in a total of 12 samples for each of the high networks, 12 for the wet root network, 11 for the wet bulk and rhizosphere soil networks, and 9 samples for each of the dry networks, as there were fewer dry sites and one wet bulk and one rhizosphere soil sample were removed due to low yield. Each subset ASV table was filtered to include only the 250 most abundant ASVs to reduce network dimensionality. Networks were inferred with SPIEC-EASI using the neighborhood selection method (MB) with 20 λ values, a λ.min.ratio of 1e–2, and a StARS instability threshold of 0.05 based on 100 subsampling iterations to ensure stability. Edges were symmetrized using the union rule, in which a connection is retained if selected for either taxon in the pair. The resulting adjacency matrices were visualized and network properties including node and edge counts, degree, betweenness, closeness centrality, modularity, transitivity, and density were computed using igraph.

The transfer of microbes from source compartments to sink compartments was modeled using fast expectation-maximizing for microbial source tracking (FEAST) (Shenhav *et al*., 2019). Simulations were run to examine the proportion of bacteria/archaea and fungi tracked from bulk soil to rhizosphere soil and from bulk and rhizosphere soil to the root endosphere for each tree sampled. Microbial transfer between each niche compartment was compared across climates with Tukey HSD tests and general linear models.

Bacterial community assembly within niche compartments and climate types was modeled with iCAMP, infer community assembly mechanisms by phylogenetic bin-based null model analysis (Ning *et al*., 2020). To limit distortions from rare taxa, 16S ASVs with less than 20 total reads across the dataset or entirely absent from more than 5 samples were removed. Remaining ASVs were aligned with the multiple sequence alignment tool, MAFFT (Katoh and Standley, 2013) using nucleotide GTR model, and the Newick format tree was built with FastTree (Price, Dehal and Arkin, 2009) in a bash environment at the default specifications. iCAMP was run with the ic.big() R function with default binning and null model settings and a full pairwise sample comparison with 999 random iterations. To analyze the iCAMP results, only within niche and within site sample pairings were considered.

## Results

### Bioinformatics results

We obtained 6,967,319 filtered reads from the 16S library and 8,642,123 reads from the ITS2 library across 99 total samples. We rarefied to 26,758 reads per sample for the entire 16S library which resulted in 2 bulk soil samples (site CR4, tree 144 and site CR1.5, tree 1708, which had 8 and 18,602 reads respectively) being removed from the dataset. For ITS, we rarefied to 23,736 reads per sample, which required removing one rhizosphere sample from CR3, tree 146. These filtering criteria effectively represent the diversity of the amplicon libraries (S6).

### Microbial diversity varies with climate

In general, as average maximum temperatures increased and elevation decreased, bulk soil alpha-diversity increased for both Bacteria/Archaea (temperature: t-val=4.943, p-value<0.001; elevation: t-val=-4.629, p-value=0.0013; Figure 2, S7) and Fungi (temperature: t-val=5.348, p-value<0.001; elevation: t-val=-5.495, p-value<0.001; Figure 2, S7). For Bacteria/Archaea, these trends were also consistent though slightly reduced in the rhizosphere (temperature: t-val=2.289, p-value=0.0478; elevation: t-val=-2.555, p-value=0.0309; Figure 2, S7), whereas for Fungi, the trends disappeared in host-associated compartments (Figure 2, S7). The only significant impact of precipitation on alpha-diversity was for root Bacteria/Archaea (t-val=2.309, p-value=0.0463; Figure 2, S7). Elevation also marginally influenced the bacterial/archaeal root alpha-diversity (t-val=-2.134, p-value=0.0617), though there were no other significant trends between these climate factors and root alpha-diversity (Figure 2, S7).

**Figure 2:**
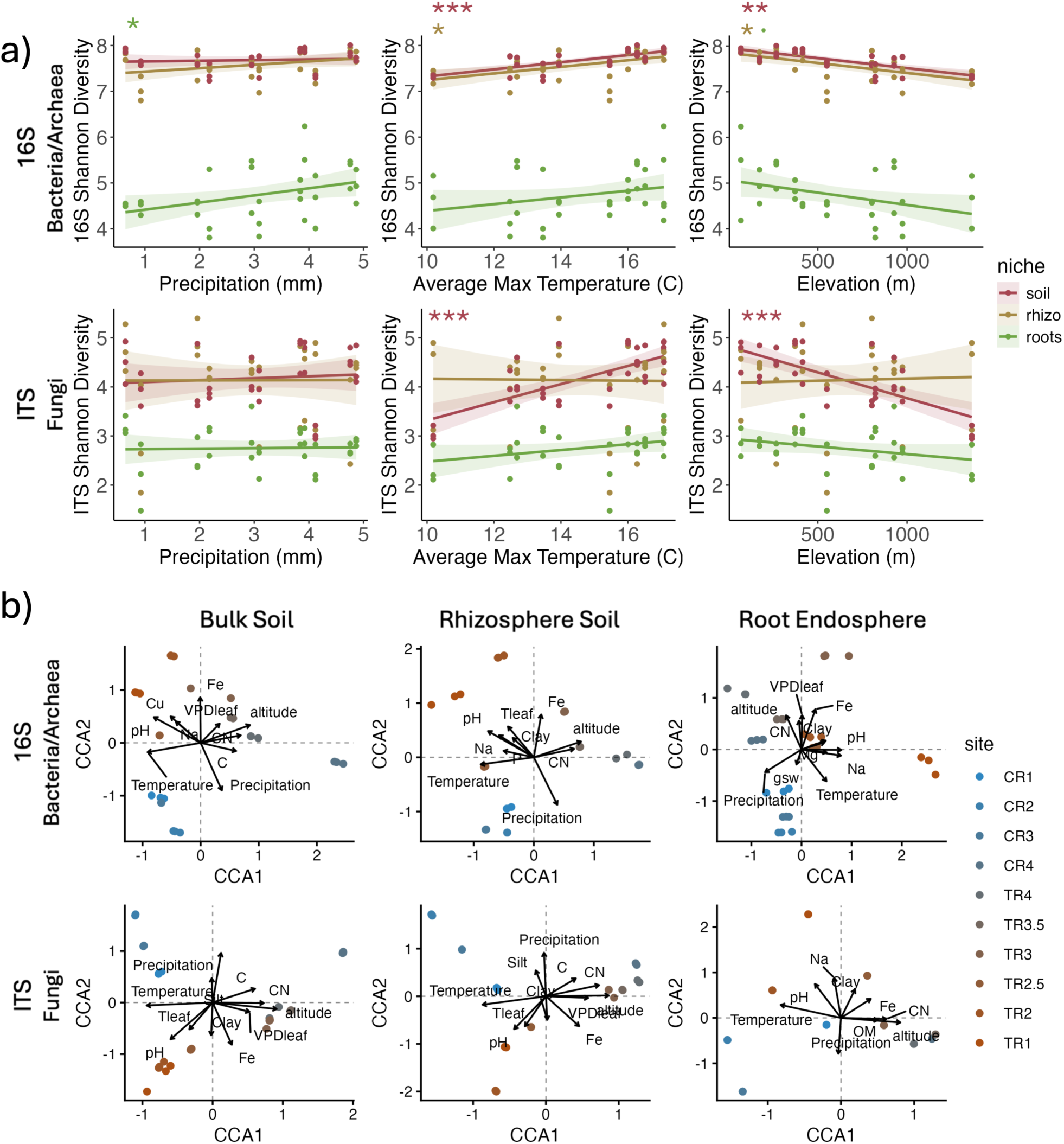
(a) Alpha-diversity trends (Shannon index) related to climate variables, average daily precipitation (left) and average daily maximum temperature (center) over the last decade, extracted from DAYMET, and elevation (right) based on site-level geographic location. Trend lines are separated and colored by niche into bulk soil (red), rhizosphere soil (yellow), and root endosphere (green). The shaded regions represent the standard deviation for each niche. Asterisks indicate significant correlations as determined by linear mixed effects models (LMEs). (b) Canonical Correspondence Analysis (CCA) plots for 16S (Bacteria/Archaea - top) and ITS (Fungi - bottom), in bulk soil, rhizosphere soil, and root endosphere (left to right) showing the influence of temperature, rainfall, altitude, soil factors (pH, Fe, CN, Na, Clay, C, Silt, Mg, Cu, K, OM, P), and host factors (VPD_leaf_, Tleaf, g_sw_). Points are color-coded by site from the west (Cowlitz River) to east (Tieton River) on a blue to red scale. CR1.5 was excluded as only shallow soil cores were collected due to rocky conditions.

For both Bacteria/Archaea and Fungi across all niche compartments, average temperature and precipitation of the last decade, altitude, and soil factors such as pH, iron content, carbon-to-nitrogen ratios (C:N), and sodium content drove beta-diversity trends (Figure 2, S8-10). Other soil factors like clay and silt content, carbon, magnesium, potassium, copper, and phosphorus, and plant photosynthetic metrics like stomatal conductance (*g_sw_*) were significant for Bacteria/Archaea or Fungi in some niches but not others (Figure 2, S8-10). For Bacteria/Archaea, temperature, altitude, and pH all had the greatest influence in the bulk soil (temperature: R^2^= 0.171, p-val<0.001; altitude: R^2^= 0.107, p-val<0.001; pH: R^2^= 0.083, p-val<0.001; S10), slightly diminished influence on the rhizosphere (temperature: R^2^= 0.159, p-val<0.001; altitude: R^2^= 0.105, p-val<0.001; pH: R^2^=0.072, p-val<0.001; S10), and a further reduction in the root endosphere (temperature: R^2^= 0.084, p-val<0.001; altitude: R^2^= 0.051, p-val<0.001; pH: R^2=^0.038, p-val<0.001; S10). Similarly to alpha-diversity, the influence of precipitation on bacterial/archaeal beta-diversity was strongest in the root endosphere (R^2^= 0.091, p-val<0.001; S10), though still significant in the rhizosphere (R^2^= 0.053, p-val<0.001) and bulk soil (R^2^= 0.064, p-val<0.001). Fungi followed slightly different trends with the effects of many variables peaking in the rhizosphere including average decadal temperature (R^2^= 0.101, p-val<0.001; S10) and soil iron content (R^2^= 0.089, p-val<0.001; S10). Of all the variables, temperature controlled the most fungal beta-diversity variance in the plant-associated compartments (rhizosphere: R^2^= 0.101, p-val<0.001; roots: R^2^= 0.080, p-val<0.001; S10) whereas for the bulk soil, fungal beta-diversity was correlated most with altitude (R^2^= 0.102, p-val<0.001; S10). Precipitation was less influential than temperature and altitude for all niches (Figure 2, S8-10). Overall, microbial beta-diversity primarily correlated with climate variability (temperature and precipitation) and secondarily with soil factors, particularly those correlating with altitude (Fe, pH, C:N; Figure 2, S8-10).

### Microbial communities cluster into three distinct climate groups

Changes in bacterial/archaeal and fungal beta-diversity were driven by site and niche (Figure 3, S11-12). When all niches (bulk soil, rhizosphere soil, and root endosphere) are evaluated together, differences in bacterial/archaeal and fungal beta-diversity were roughly differentiated by the elevation gradient with considerable overlap between the low-elevation sites on the Tieton and Cowlitz watersheds (S11). Interestingly, bacterial/archaeal beta-diversity was driven to a greater extent by niche (F-stat=18.300, p-val<0.001; S12) than by site (F-stat=4.562, p-val<0.001; S12) whereas for fungi, the opposite trend occurred (Niche: 3.674, p-val<0.001, Site: 4.025, p-val<0.001; S12), though fungal beta-diversity differences were less than bacterial/archaeal overall (Bacterial/Archaeal Residuals: R^2^=0.354, Fungal Residuals: R^2^=0.466; S12).

**Figure 3:**
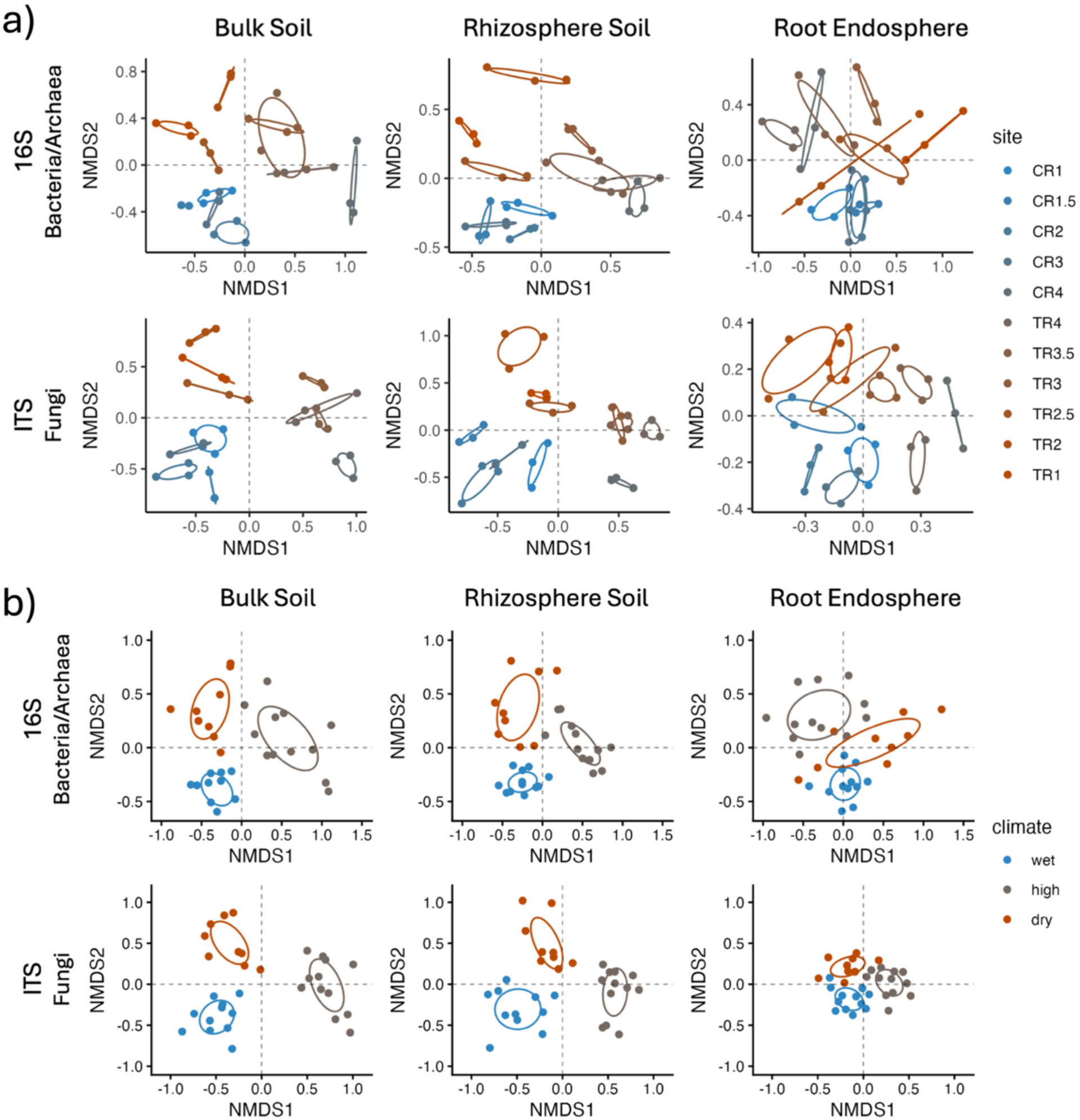
(a) Non-metric multi-dimensional scaling (NMDS) plots showing colored by sites on a blue to orange scale for bulk soil (left), rhizosphere soil (center), and root endosphere (right) for bacterial/archaeal beta-diversity (top row) and fungal beta-diversity (bottom row). Beta-diversity was calculated based on Bray-Curtis dissimilarities. (b) NMDS ordinations clustered and color-coded by climate category. Blue points represent the wetter sites on the lower Cowlitz watershed, grey represent the cooler, high altitude sites at the top of the Cowlitz watershed and the upper Tieton watershed, and the red points are the drier sites on the lower Tieton watershed.

Within niches, bacterial/archaeal and fungal beta-diversity was driven by site, and site differences were slightly diminished in host-associated compartments, with the greatest site differences in the bulk soil, and reduced effects in the roots for both Bacteria/Archaea and Fungi (Figure 3, S11). When separated by niche compartment, samples formed rough but consistent clusters by climate. The four lowest Cowlitz watershed sites formed a “wet” cluster, the four highest elevation sites formed a “high” group, and the three lowest Tieton watershed sites formed a “dry” group (Figure 3). When all niches were considered together, there was some overlap between the wet and dry categories for both Bacteria/Archaea and Fungi (S11) suggesting that beta-diversity was driven more by elevation/temperature than precipitation. When niches were considered separately there was stronger clustering for the soil environments, while the root endosphere had slightly reduced though still significant beta-diversity differentiation by climate category for both Fungi and Bacteria/Archaea (Figure 3, S11).

### Trees show signs of stress in high and dry sites compared to wet sites

g_sw_, VPD_leaf_, and ΦPSII all vary along the transect (S13), driven by varying temperature and precipitation, soil factors, and elevation (S14-15). g_sw_ correlated with precipitation (r=-0.139, p-val=0.043; S15) but not with temperature or altitude, indicating that stomatal closure was linked most closely to precipitation-moisture related conditions. VPD_leaf_ increased with reduced precipitation (r=-0.612, p-val<0.001; S14-15) and peaked at intermediate elevation (r=0.329, p-val=0.001; S14-15) and overall was lower in the wet climate than in the high and dry (Figure 4). ΦPSII increased with precipitation (r=0.371, p-val<0.001; S14-15) and temperature (r=0.297, p-val<0.001; S14-15) and reduced with altitude (r=-0.379, p-val<0.001; S14-15). It was slightly lower and noteably more variable in the high and dry climate than in the wet climate (Figure 4). g_sw_, VPD_leaf_ and ΦPSII all varied considerably more across sites and climates than from tree-to-tree within sites (S17). Overall, trees within wet sites faced less photosynthetic stress than in high and dry sites.

**Figure 4:**
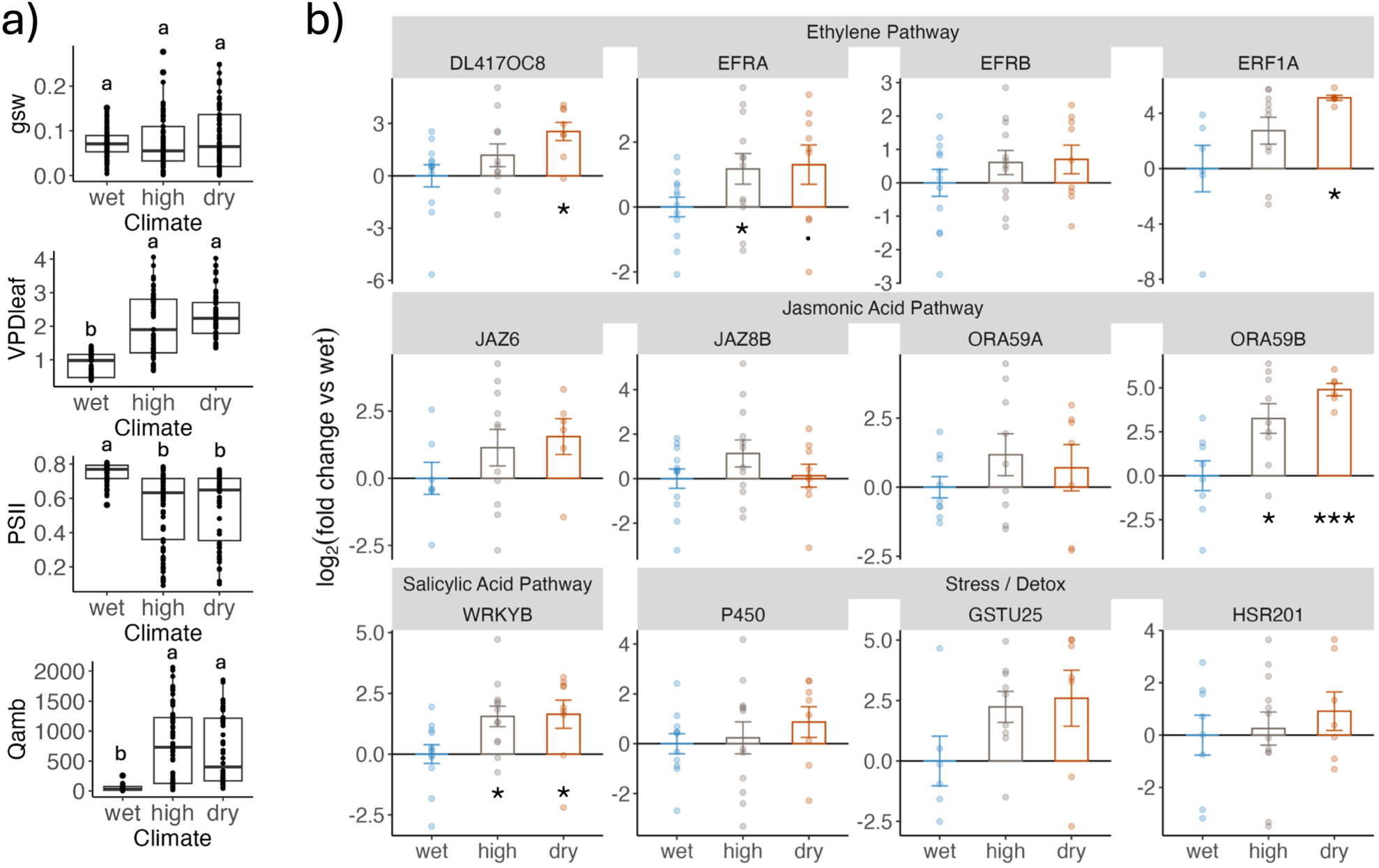
(a) Tree ecophysiological data for (top to bottom) stomatal conductance to water vapor in mol m^-2^ s^-1^ (g_sw_), vapor pressure deficit at the temperature of the leaf in kPa (VPD_leaf_), photosystem II in 1 – steady state flux / maximum flux (PSII), and ambient light in µmol of photons m⁻² s⁻¹ (Qamb) by climate category. Letters indicate Tukey HSD groups. (b) Quantitative Reverse Transcription Polymerase Chain Reaction (qRT-PCR) results for known stress response genes in *Populus trichocarpa* foliar samples. UBQ10 and 18S were used as housekeeping genes for normalization and the average gene expression in the wet sites was used as a treatment control. Higher log2fold change compared to the wet site indicates increased gene expression. Asterisks (*) indicate significant differences in gene expression compared to the wet climate category as determined with student t-tests.

The expression of abiotic stress genes involved in ethylene, jasmonic acid and salicylic acid pathway and 3 genes involved in general abiotic stress response and detoxification (S18) all increased in the high and dry sites compared to the average expression across the wet sites, typically with the greatest expression in dry sites (Figure 4, S19). The ethylene pathway genes tested––DL4170C8, EFRA, EFRB, and ERF1A––increased most in dry sites (Figure 4), significantly for DL4170C8 (ddCq=-2.541, p-val=0.011, S19) and ERF1A (ddCq=-5.115, p-val=0.013, S19) and marginally significant in EFRA (ddCq=-1.305, p-val=0.052, S19). EFRA also significantly increased at high-altitude compared to wet sites (ddCq=-1.173, p-val=0.048, S19), though the other ethylene pathway genes were variable and expression differences were otherwise non-significant (Figure 4, S19). Jasmonic acid pathway gene expression––JAZ8B, JAZ6, ORA59A, and ORA59B––increased slightly in high and dry compared to wet, where some increased most in dry (JAZ6: ddCq=-1.555; ORA59B: ddCq=-4.294, S19) and others in high-altitude sites (JAZ8B: ddCq=-1.136; ORA59A: ddCq=-1.190, S19). Most of the jasmonic acid pathway genes showed non-significant trends, except for ORA59B which significantly increased in both high-altitude (p-val=0.035, S19) and dry sites (p-val<0.001, S19) compared to wet. WRKYB, a salicylic acid pathway gene, significantly increased expression in both high-altitude (ddCq=-1.552, p-val=0.012, S19) and dry sites (ddCq=-1.639, p-val=0.024, S19) compared to wet sites. For many genes, expression varied most in the high-altitude category, especially for jasmonic acid pathway genes (Figure 4, S19).

The relative expression of some genes also correlated to fungal and bacterial/archaeal beta-diversity (S20-21). ORA59B, correlated with both fungal (F-stat=2.069, p-val=0.001) and bacterial/archaeal beta-diversity (F-stat=0.014, p-val=0.006; S21). JAZ8B, another jasmonic acid response gene, and WRKYB, a gene on the salicylic acid response pathway were both correlated with fungal communities across niches (JAZ8B: F-stat=1.680, p-val=0.003; WRKYB: F-stat=1.533, p-val=0.015, S21). For many genes sampled, the variability in gene expression within climate types was greatest for the high-altitude category. This trend was most apparent for the jasmonic acid pathway genes (Figure 4, S19). However, there were no significant correlations between the root microbial communities and the foliar stress response (S21).

### Network structure and microbial co-occurrence patterns vary with niche and climate category

Across all climate types, there were more fungal nodes than bacterial in both the bulk and rhizosphere soil, though the trend reversed in the root endosphere (Figure 5, S22). In the wet climate, the number of edges increased from bulk to rhizosphere, to root, indicating an increase in co-occurring taxa in host-associated environments (Figure 5, S22). Dry networks followed a similar trend with the root network having the greatest number of edges, though the bulk and rhizosphere were more comparable (Figure 5, S22). Interestingly, this trend reversed in the high networks, where bulk had more edges than the rhizosphere, and the root had the fewest (Figure 5, S22). High networks had slightly greater modularity and transitivity (S22). Network density was slightly higher in the high and dry climate types (S22). Network metrics also varied with niche compartment. Modularity reduced in host-associated niches in the wet and dry networks but increased in high networks (S22). Transitivity was higher in the bulk soil in the wet and dry climate networks but spiked in the roots in the high networks (S22). This observation is possibly driven by a few highly connected fungal modules (Figure 5). Visually, the networks appear similar by climate class. Most networks had a main cluster of highly central bacterial and fungal ASVs. The stressed climate networks often had secondary modules that were almost exclusively fungal, populated primarily by Ascomycota (Figure 5). In the high-elevation root network, there were a few modules of highly co-occurring Ascomycota and Basidiomycota that had few connections with the other network nodes (Figure 5). While fungal nodes outnumbered bacterial nodes in the bulk soil and rhizosphere networks of every climate type, the taxa with the greatest degree, closeness centrality, and betweenness were a mix of Fungi and Bacteria (Figure 5, S23). A few members of Actinomycota and Ascomycota were highly central and connected across all networks, but especially in the dry networks for every sample type (Figure 5, S23). Members of Pseudomonadota were highly central and connected in the wet and high climate networks but not in the dry networks (Figure 5, S23), which aligns with relative abundance trends by niche and climate (S24-25).

**Figure 5:**
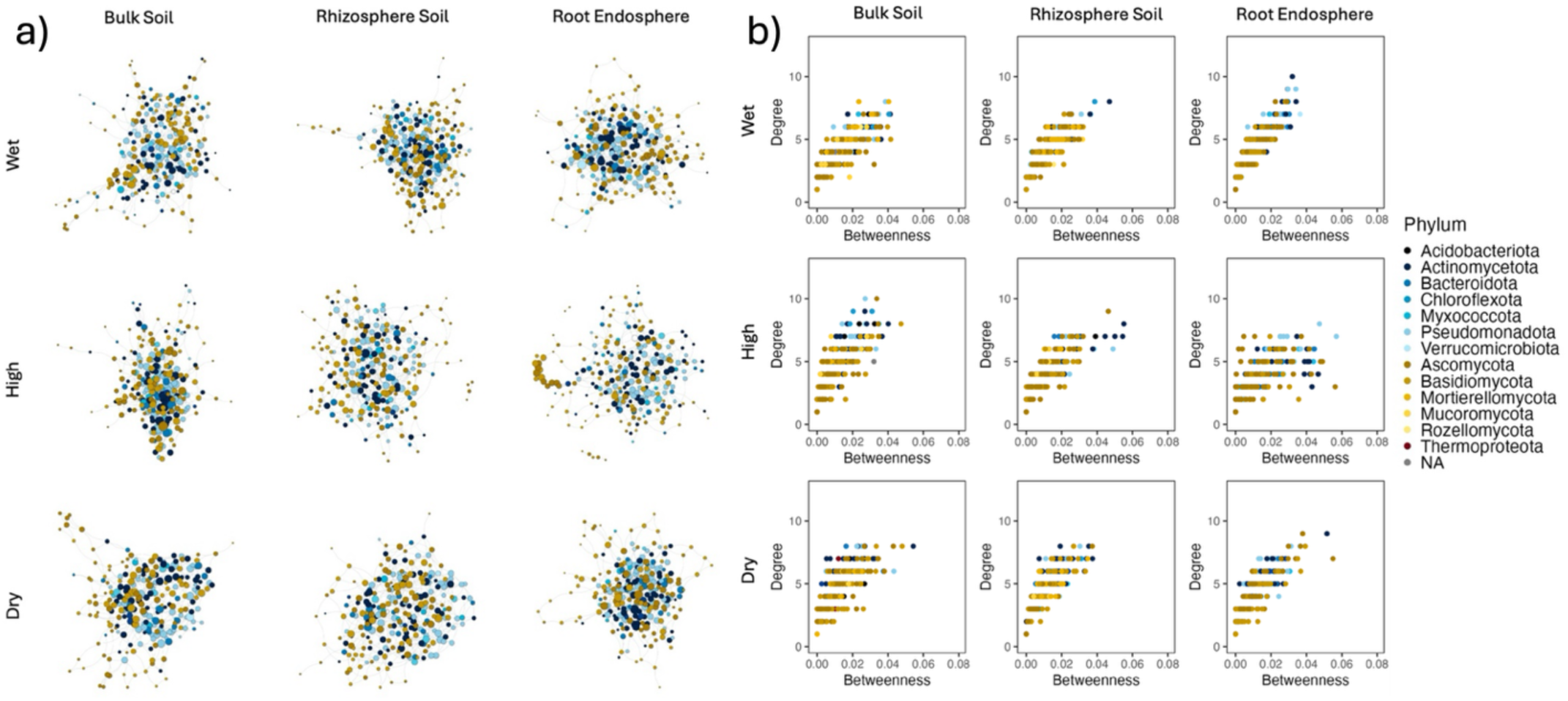
(a) Co-occurrence networks generated with the neighborhood selection (MB) method in SpiecEasi and visualized with igraph, showing the inferred associations among bacterial ASVs (blue nodes), fungal ASVs (yellow nodes), and archaeal ASVs (red nodes). Edge presence indicates a non-zero conditional association inferred by the MB method. The size of the node is scaled by its degree (number of edges connecting it to other nodes). Samples were divided by niche (left to right: bulk soil, rhizosphere soil, and root endosphere), and climate classification (top to bottom: wet, high sites, and dry). (b) Graphs showing the degree (number of edges) vs. the betweenness of each node within the network. Every point represents a node within the network and is color-coded by phylum.

### Roots in wet sites sourced a greater proportion of soil Fungi than roots in high-altitude and dry sites

The transfer of microbes from the bulk soil community to the rhizosphere was slightly higher under stress conditions (Figure 6). In the wet rhizospheres, an average of 75.**7**% of Bacteria and 81.7% of Fungi were sourced from the bulk soil community, which was slightly lower than in the high sites where 77.2% of Bacteria and 82.**9**% of Fungi were sourced from the bulk soil (Figure 6). Dry sites had the highest proportional transfer from bulk to rhizosphere soil with 80.4% of Bacteria and 83.8% of Fungi. The root endosphere community was primarily sourced from unknown pools, except for the root fungal communities in wet sites, which sourced an average of 55.**3**% from the rhizosphere and 4.6% from the bulk soil. For both Bacteria and Fungi, the prevalence of microbes from unknown sources in the root endosphere was higher in the stressed climates (Figure 6). On average, roots in high sites sourced 68.**2**% of their Bacteria and 64.**4**% of Fungi from unknown sources, while roots in dry sites sourced 64.**4**% of Bacteria and 64.3% of Fungi from unknowns (Figure 6). Fungi had a steeper increase in the proportion of root organisms sourced from unknown pools in high and dry compared to wet than Bacteria, which could indicate that Fungi are more responsive to environmental conditions. Tree-to-tree variability was also higher in Fungi, which could be caused by different host and fungal responses to slight soil and climate differences within each climate class, or due to greater heterogeneity of fungal communities than bacterial from site to site.

**Figure 6:**
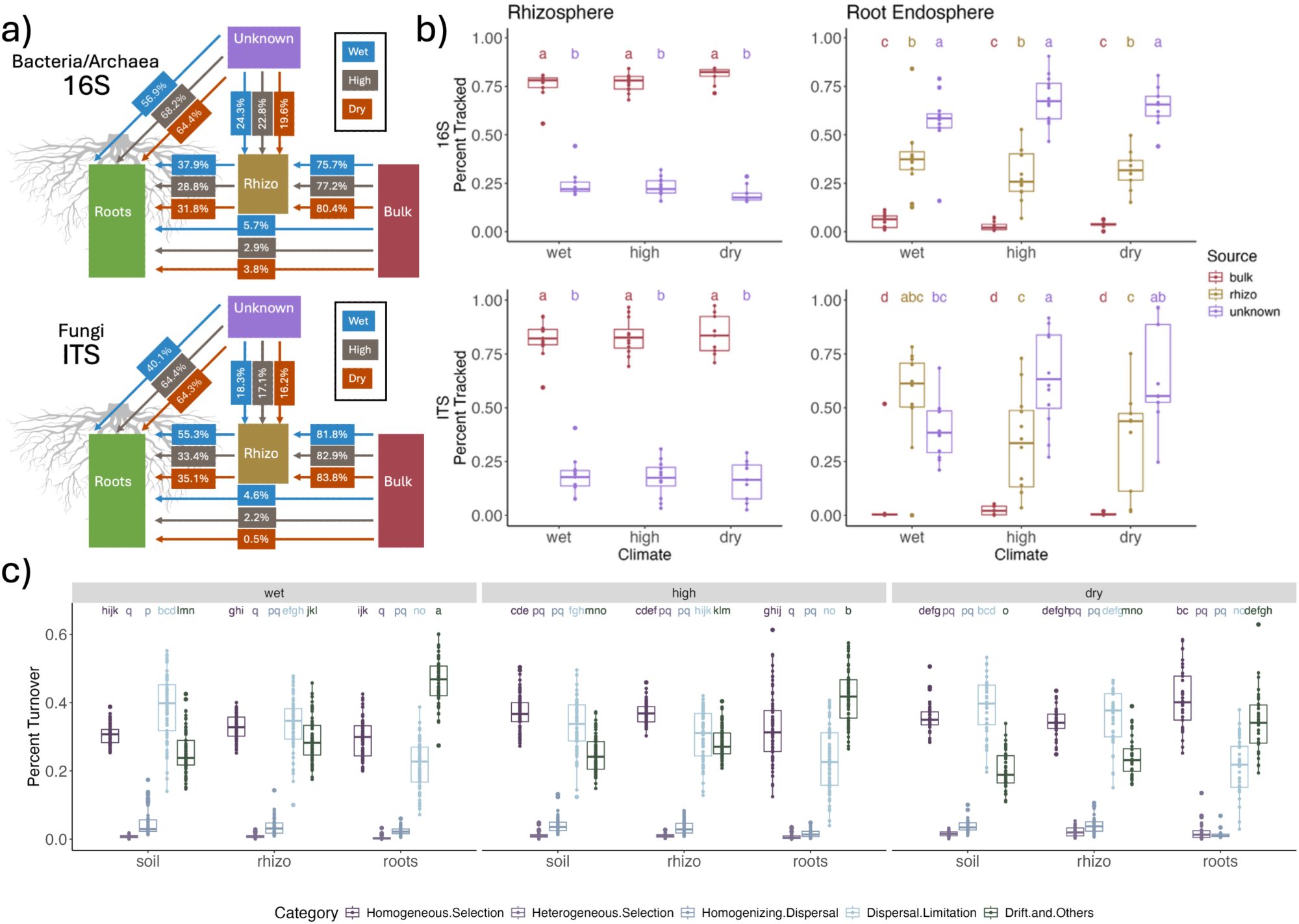
(a) FEAST Source Tracker analysis of microbial transfer from bulk soil to rhizosphere soil and from bulk and rhizosphere soil to the root endosphere for each tree sampled. Flow charts show the average percent transfer in the wet, high, and dry treatments. (b) Boxplots with Tukey HSD significance labels tracking from the source (bulk soil in red, rhizosphere soil in yellow, and unknown sources in purple) to the rhizosphere soil and the root endosphere. Source tracking was calculated for 16S (top) and ITS (bottom). (c) iCAMP modeling of the 16S amplicon library to determine drivers of microbial community turnover. Selection processes like homogenous selection (dark purple) and heterogeneous selection (light purple) and stochastic processes like homogenizing dispersal (dark blue), dispersal limitation (light blue), and drift and others (dark green) were all considered. Only within niche and within climate sample comparisons were considered. Letters represent Tukey HSD groups.

### Bacterial community turnover in the root endosphere was driven by selection processes in the dry climate but not in the wet and high climate

Overall, selection processes had a greater influence on bulk and rhizosphere soil bacterial community turnover in high-altitude and dry sites than in wet sites (Figure 6). The impact of selection on bacterial communities was higher in the root endosphere (43.3%) compared to the bulk (37.2%) and rhizosphere soil (36.5%) in the dry climate, though in the wet climate, selection processes functioned similarly across compartments (bulk=31.5%, rhizo=33.8%, root=29.9%), and in the high climate selection actually decreased in the roots (bulk=38.2%, rhizo=37.9%, root=33.4%; Figure 6, S26). Total stochastic processes were higher in the roots of the high-altitude (66.6%) and wet sites (70.1%) compared to their respective rhizosphere soils (high=62.1%, wet=66.2%), whereas in the dry sites, stochastic processes did not increase in the roots (bulk=62.8%, rhizo=63.5%, root=56.7%; Figure 6, S26). Heterogeneous selection was a minor influence overall, but slightly greater in the dry climate across all niche compartments (Figure 6, S26). Homogenous selection was a strong influence overall and was a greater driver of turnover in the stressed, high-altitude and dry climates than in the wet climate. Of the stochastic processes, dispersal limitation had the greatest influence on bulk and rhizosphere soil across all climates but was not as major a driver of turnover within the root endosphere, where drift and others dominated the stochastic category (Figure 6, S26).

## Discussion

### Alpha-diversity trends differed by microbial kingdom and niche compartment

Bacterial/archaeal bulk soil and rhizosphere alpha-diversity correlated with temperature and was inversely correlated with altitude, indicating that cold temperatures or low soil nutrient status associated with high elevations limited bacterial/archaeal diversity (Figure 2), aligning with the majority of gradient studies (Wang *et al*., 2022). Fungal diversity was also inversely correlated with elevation in our study (Figure 2), however fungal trends in the literature are more variable with many studies finding opposite trends (Shen *et al*., 2020; Ma *et al*., 2022). Microbial communities are dependent on a variety of factors, and bacterial, archaeal and fungal diversity changes with elevation are likely a reflection of temperature and soil variables, like pH (Shen *et al*., 2019). While bacterial/archaeal diversity of the rhizosphere and root endosphere followed the same trends as the bulk soil, we did not observe significant patterns for rhizosphere or root endospheric fungi (Figure 1). Soil pH is generally inversely correlated with elevation and is a strong driver of microbial diversity both in our study (Figure 2) and others (Wang *et al*., 2015; Shen *et al*., 2013; Zhang *et al*., 2015). pH often has a slightly stronger influence on bacterial/archaeal communities than fungal communities, as Fungi generally tolerate broader pH ranges (Rousk *et al*., 2010). Many of these alpha-diversity trends may be system specific, with factors like vegetative cover, aspect of exposure, climate type, and studied spans of elevation leading to different observations.

### Climate influenced microbial beta-diversity more than soil variables

Temperature, precipitation, and elevation were all strongly correlated with bacterial/archaeal and fungal beta-diversity in all niche compartments in our study (Figure 2, S10), which is surprising as soil factors like pH, soil nutrients, and soil physical properties are often more influential than climate factors like temperature on the soil microbiome (Wang *et al*., 2022). In our case, pH is correlated with altitude and temperature, likely due to the shared effects of the altitude gradient, however the correlations between the microbial communities and altitude and temperature were still stronger than correlations with soil factors when tested independently (S9). For both bacterial/archaeal and fungal communities, the effects of temperature and altitude were greater than those of any single soil factor, though pH had a greater individual influence on bacterial/archaeal communities than precipitation, and C:N ratios and iron content played a larger individual role than precipitation for fungal communities (S9). Our results indicate that climate is driving compositional changes in both soil and plant-associated environments.

### The climate gradient drives changes in bulk soil, rhizosphere soil, and root endophytic microbial communities

The environmental conditions along the transect created three distinct microbial communities: a wet community along the Cowlitz River, a cool and high community in the highest elevation sites, and a dry community in the lower elevation sites along the Tieton River in a variety of analyses (Figure 3). The distances between climate groups were slightly greater for bulk and rhizosphere soil Fungi than Bacteria/Archaea (Figure 3), possibly due to greater spatial heterogeneity and distribution limitation of Fungi, compared to Bacteria which tend to be slightly more spatially homogenous and dispersible (Štursová *et al*., 2016; Bazany *et al*., 2022). However, bacterial/archaeal communities are also distinct by climate, which could be driven by different conditions providing advantages to different taxa. For example, dry sites likely favor organisms capable of dormancy and sporulation (Coleine *et al*., 2024). In both bacterial/archaeal and fungal communities, oligotrophic phyla were broadly enriched in the dry sites, for example Actinomycetota and Chloroflexota Bacteria and Ascomycota Fungi (Figure 5, S21, S24-25).

As this is a natural system with mature trees, we are likely observing fairly persistent, as opposed to transitional communities. In short-term osmotic stress, microbial transcription shifts, and some taxa undergo dormancy or other adaptive measures, whereas in longer time-scales microbial enrichment patterns change (Coleine *et al*., 2024; Bazany *et al.,* 2025). In a broad sense, bacteria and fungi respond differently to short- and long-term osmotic stress, where fungi are more resistant, with community and network architecture taking longer to shift under drought conditions, and bacteria are more resilient, recovering to return to pre-drought community structure faster than fungi (de Vries *et al*., 2018; Gao *et al*., 2022).

### Elevation and dry conditions result in similar physiological stress in *Populus trichocarpa* but more variable expression of key stress genes

*P. trichocarpa* trees in our study displayed similar g_sw_, VPD_leaf_, and ΦPSII in dry and high sites compared to wet sites (Figure 4). This could be driven by higher ambient light in high and dry climates (Figure 4), as transpiration and photosynthetic processes can be influenced by radiation damage or shading effects (Körner, Allison and Hilscher, 1983; Tyagi *et al.,* 2016). Changes in tree physiology could also be driven by soil factors, as higher altitudes generally have lower soil nutrients due to slower decomposition rates, leaching, and aridity. Tree genotype differences could also influence plant stress phenotypes (Byars, Papst, Hoffmann, 2007). Tree genotypes may also vary along the transect (Zhang, Suren and Holliday, 2019; Kooyers, 2015), and localized adaptations to aridity and high-altitude stress may have homogenized the responses to different stressors.

Although the physiological stress was similar in high and dry sites, there were differences in the gene expression of foliar stress indicators. For a panel of twelve genes within the ethylene, jasmonic acid, salicylic acid, and other stress and detox pathways, relative expression increased in the foliar tissue in high and dry sites compared to the wet “control” sites (Figure 4, S19). Ethylene pathway genes are often enriched under abiotic stress (Achard *et al*., 2006). Ethylene synthesis is involved in regulating stomatal closure, root growth and leaf senescence, among other processes under drought (Arraes *et al*., 2015; Fatma *et al*., 2022), and high altitude can also drive changes to ethylene response pathways (Vall-llaura *et al*., 2025). While cooler temperatures associated with higher altitudes can result in reduced ethylene accumulation, many Angiosperms respond to the reduction in partial pressure of oxygen in part by altering the transcription of certain ethylene response factors (Abbas *et al*., 2022). ERF genes are often upregulated in response to reactive oxygen species accumulation under cold stress in order to increase antioxidant enzyme activity (Lee *et al*., 2020).

Genes related to jasmonic acid synthesis and detection are also frequently upregulated in plants under many types of abiotic stress including osmotic stress. Jasmonic acid genes and ethylene genes are often in dialogue, sometimes inhibiting one another and other times acting synergistically (Yamamoto *et al*., 2020). During drought, jasmonic acid accumulation can initiate stomatal closure as well as the production of various osmoprotectants like proline and soluble sugars (Xing *et al*., 2020). Jasmonic acid also interacts with abscisic acid in *Populus* species, an important signal in osmotic stress regulation (Rao *et al*., 2023). Stresses associated with high altitude like increased solar radiation and freezing can also be alleviated by jasmonates through many of the same mechanisms active in alleviating drought stress (Ali and Baek, 2020) as many of the molecules synthesized through jasmonic acid response pathways that act as osmoprotectants under desiccation stress can also function as cryoprotectants under cold conditions (Li *et al*., 2024).

WRKYB, a salicylic acid response gene, had increased expression in both high altitude and dry sites. Salicylic acid can stimulate the production of osmoprotectants like proline (La *et al*., 2019). Both salicylic acid and ethylene interact with dehydration-responsive element-binding proteins, key transcription regulators involved in drought stress response (Han *et al*., 2022). Salicylic acid genes also respond to stresses like cold temperatures and increased UV radiation (Zhang, Tonsor and Traw, 2015), which could explain the increased expression of WRKYB at high-altitude sites.

The more general stress response and detox genes were all upregulated in high and dry sites compared to wet sites, however the trends were quite variable. This could be because they are more responsive to higher levels of abiotic stress, and the caliber of stress varies across the transect, or because the pathways the genes impact are quite variable. P450 binds oxygen to hydroxylate flavonoids, an intermediate step in the formation of secondary pigments and antioxidant flavonols (Azaiez *et al*., 2009). GSTU25 is a stress-responsive class of glutathione S-transferase, an enzyme that defends the plant against oxidative stress (Lin *et al*., 2025). HSR201 is a gene involved in the hypersensitive response pathway, triggering cell death and salicylic acid production in response to a variety of biotic and abiotic stress indicators including fungal and abiotic elicitors (Guo and Cheng, 2022).

While the expression of a few of the stress-response genes in the foliar tissues correlated with the overall microbial beta-diversity (S20-21, these trends diminished in the root endosphere (S20-21), indicating that while both fungal beta-diversity and the expression of certain stress-response genes were influenced by climate, the expression of the genes in this panel were not a primary driver of root community composition. It should also be noted that in *Populus*, an evolutionarily recent genome duplication event has resulted in highly tissue-specific expression patterns for many highly similar, but paralogous gene copies (Segerman, Jansson and Karlsson, 2007). Such tissue-specific expression patterns could also have contributed to the observed lack of correlations between foliar gene expression and root microbial communities.

### Patterns of microbial cooccurrence varied with climate type and niche compartment

Network architecture varied slightly by climate type and niche compartment (Figure 5). Overall, bulk soil networks were more connected under both high and dry conditions than root endosphere networks, with a greater number of edges and higher network densities (Figure 5, S22). Rhizosphere networks were also robust to stress with relatively high density (S22). Compared to the bulk soil networks, the rhizosphere networks had higher transitivity (S22), which may be evidence of stronger, localized interactions between bacterial and fungal taxa in response to plant exudation. Root networks were slightly less robust to the high and dry climate, with fewer edges and generally lower density compared to the bulk and rhizosphere soils (S22). In general, root endosphere networks, were sparse and modular (Figure 5), consistent with top-down filtering by environmental conditions and host selection. Interestingly, wet networks had slightly lower density and fewer edges than dry and high networks, contrary to what we often see in drought studies (Bazany *et al*., 2022; Gao *et al*., 2022), which could be an indication that high-altitude and microbiome communities under more consistent arid conditions, particularly those in the soil, have stabilized over time. This likely reflects long-term adaptation in these naturally dry or high-elevation systems rather than acute stress effects typically seen in experimental drought studies.

The dominant microbial hubs varied with climate. Members of the Ascomycota phylum were frequently identified as hubs based on high degree, closeness centrality, and betweenness, under both the high elevation and the dry stress conditions (Figure 5, S22-23). We observed increases in Ascomycota and Basidiomycota and a depletion of Mortierellomycota and Mucoromycota in the root endosphere in each site and climate category (S24-25). These fungal modules are often connected to common bacterial/archaeal nodes which could indicate the presence of mycorrhizal fungi and mycorrhizae helper bacteria which facilitate the establishment of mycorrhizal fungi in and around plant roots (Sangwan and Prasanna, 2022) or negative correlations which could indicate antagonism between taxa. Pseudomonadota were frequent hubs in wet and high-elevation networks, particularly in the roots, however they were not prominent in any of the dry networks (Figure 5, S23). While some members of the Actinomycetota phylum were identified as hub taxa across the climate types and niche compartments, they were the most prominent bacterial hubs in the dry networks (Figure 5, S23). In arid conditions, plant immune responses are often suppressed in favor of basic survival mechanisms (Bostock, Pye and Roubtsova, 2014; Castrillo *et al*., 2017; Finkel *et al*., 2019). In these cases, trees may outsource defense processes to bacterial endophytes like Actinomycetes with their broad classes of anti-biotic compounds (Van der Meij *et al.,* 2017), a phylum that is broadly enriched under drought stress (Naylor *et al*., 2017; Xu *et al*., 2018) and their induction of plant defense pathways, priming the immune system against future colonization (Conn, Walker and Franco, 2008).

### More Fungi from unknown sources colonize the root endospheres of high and dry trees

Generally, root endosphere microbes are sourced from the surrounding soil, where microbes that can pass plant physical and chemical barriers can colonize the root endosphere (Trivedi *et al*., 2020). However, some microbes can be sourced from other reservoirs, for example parent to offspring transfer (Frank, Saldierna Guzmán and Shay, 2017) as clonal reproduction by branch abscission and breakage as well as root initiation is quite common in *Populus* spp. (Brattne *et al.,* 1996). In the aboveground compartments, wind transfer and deposition can be a major factor (Ottesen *et al*., 2016), especially in dry climates where organisms are more likely to sporulate and dust is more likely to become airborne, though surrounding soil is a major source of aerial microbes (Zhou *et al*., 2021). In this study, trees accumulated a higher proportion of their root endosphere community from the surrounding soil in wet sites than in high or dry sites (Figure 6). This trend was particularly notable for Fungi, though present for the bacterial communities as well. This could be because under stressful conditions, fungal organisms found in undetectable abundances in the soil bank, for example as resistant spores, proliferate in the root endosphere.

The Fungi found in the roots that were not found in the rhizosphere and bulk soil were a mix of known endophytes, including dark septate endophytes, and organisms along the saprophyte – mutualists spectrum. Several *Mycena spp.*, an *Atractospora spp.* and a *Cyclocybe spp.* appeared across climate types (S26) which are all common in healthy *Populus* roots (Gottel *et al*., 2011; Shakya *et al*., 2013; Cregger *et al*., 2018). *Mycena spp.* occupy broad roles across the saprophyte-mutualist continuum (Thoen *et al*., 2020; Harder *et al*., 2023). Interestingly, the three dark septate endophytes, two *Cladophialophora spp.* and *Phialocephala fortinii* were all found in high-altitude roots though one of the *Cladophialophora spp.* was also found in the wet climate (S26). Dark septate endophytes can help plants at high altitude by helping the plant to acquire nutrients to alleviate stress caused by low nutrient soils (Schadt, Mullen and Schmidt, 2001; Schmidt *et al*., 2008; Malicka, Magurno and Piotrowska-Seget, 2022) as well as inducing the plant to accumulate soluble sugars, prolines, and other molecules which can protect the roots from either drought or cold (Lu *et al*., 2025; Roychowdhury *et al*., no date).

*P. trichocarpa* exerts greater selection on root bacterial communities in low precipitation sites but not in high or wet sites.

Overall, we found that bacterial turnover due to selection is higher in high altitude and low precipitation sites than in high precipitation sites, indicating that the bacterial communities are impacted in a top-down manner by both stresses associated with high-altitude and dry conditions (Figure 6) (Simonsen, 2022). There is an additional increase in the prevalence of selection processes in the root endosphere compartment compared to the soil environment in the dry, low precipitation sites, but this trend is not present in either the high or wet climate (Figure 6). This could indicate that under osmotic stress, the plant is exerting additional selective pressure on its root endosphere bacterial community, though altered root exudation profiles (Sharma *et al*., 2025) and reductions in immune system function that allow more microbes to colonize the root (Fitzpatrick *et al*., 2026). The absence of increased selection in the root endosphere bacterial community under high-elevation stress may be because there are fewer notable plant-bacterial partnerships to address stresses associated with high elevation compared to dry condition adaptation. Overall, the combined source tracking and bin-based null model analysis may indicate that *P. trichocarpa* has primarily fungal-derived microbial methods for addressing high-altitude stress and both bacterial and fungal-based methods for dealing with osmotic stress.

### Conclusions

We examined the bulk soil, rhizosphere soil, and root endosphere microbiomes of *Populus trichocarpa* over a naturally occurring climate gradient and found that microbiomes varied continuously with climate and soil factors, and categorically by niche compartment, and broad climate classes. While plants are stressed by both high-altitude and low-precipitation conditions, specific gene expression varies, with some indicator genes showing the greatest upregulation in high-altitude and others in the dry climate. We also found that microbiome communities were less divergent across the climate gradient in the root endosphere than the rhizosphere and bulk soil, indicating that while there is some consistency with host selection across environmental contexts, the climatic and other abiotic conditions, like soil properties, still drive major changes in belowground microbial communities in both soil and plant-associated environments. Thus, our study provides valuable insights into the complex relationships between climate, soil, host, and microbiome in naturally occurring environmental gradients, illustrating the importance of large-scale field studies in elucidating the factors dictating microbial community structure under variable environmental conditions.

## Acknowledgements

The authors wish to thank the Gifford Pinchot National Forest, Okanogan-Wenatchee National Forest, Washington State Department of Fish and Wildlife, Washington State Department of Natural Resources, and Tacoma Power and Light for providing access to research sites. We also thank the Cayuse, Chehalis, Klickitat, Cowlitz, Siletz, Umatilla, Walla Walla and Yakama peoples on whose ceded and unceded lands the research took place.

This research was sponsored by the Genomic Science Program, United States Department of Energy, Office of Science, Biological and Environmental Research, as part of the Plant-Microbe Interfaces Science Focus Area (https://pmiweb.ornl.gov/) at Oak Ridge National Laboratory (ORNL). ORNL is managed by UT-Battelle, LLC, for the United States Department of Energy under contract DEAC05-00OR22725.

## Data availability

All sequencing read data, soil chemistry data, tree phenotyping data, and qRT-PCR data are publicly available (DOI: 10.25983/PMI/3001412) and can be accessed through or through OSTI (ID: 3001412) at: https://www.osti.gov/biblio/3001412.

Sequencing read data is also available on NCBI SRA, PRJNA1514744.

All climate data was collected from DAYMET:

Thornton, P.E. *et al*. (2014) *Daymet: Daily Surface Weather Data on a 1-km Grid for North America, Version 2.* Oak Ridge National Lab. (ORNL), Oak Ridge, TN (United States). Available at: https://www.osti.gov/biblio/1148868 (Accessed: February 17, 2025).

R-scripts for all bioinformatics, statistical analysis, and data visualizations are available on github: https://github.com/kbazany/washington_transect_amplicon

## Supplemental Information

**S1:** Temperature and soil water values were extracted from the Terraclimate gridded dataset (Abatzoglou *et al*., 2018). Site coordinates over the Cascades in Washington State, USA, were matched to the nearest grid cell by minimizing Euclidean distance in latitude and longitude. Stress gradients were computed for each season (winter, spring, summer, fall) separately. Seasonal mean maximum temperature (°C) and soil water content (mm) were averaged over the past 10 years (January 2010 – December 2019) and normalized to unitless values between 0 and 1, where higher normalized temperature represented thermal stress, and lower normalized soil water content represented hydrological stress. A combined “stress score” was computed using the mean of the temperature and soil water components, yielding values between 0 (low stress) and 1 (high stress), where higher scores indicated hotter and/or drier conditions (Lagergren *et al*., 2023). From these options, final sites were chosen according to safety considerations, accessibility, permit procurement, broad soil characteristics, and the number of sampleable *P. trichocarpa* trees, with a minimum of 3 trees per site required.

**S2:**
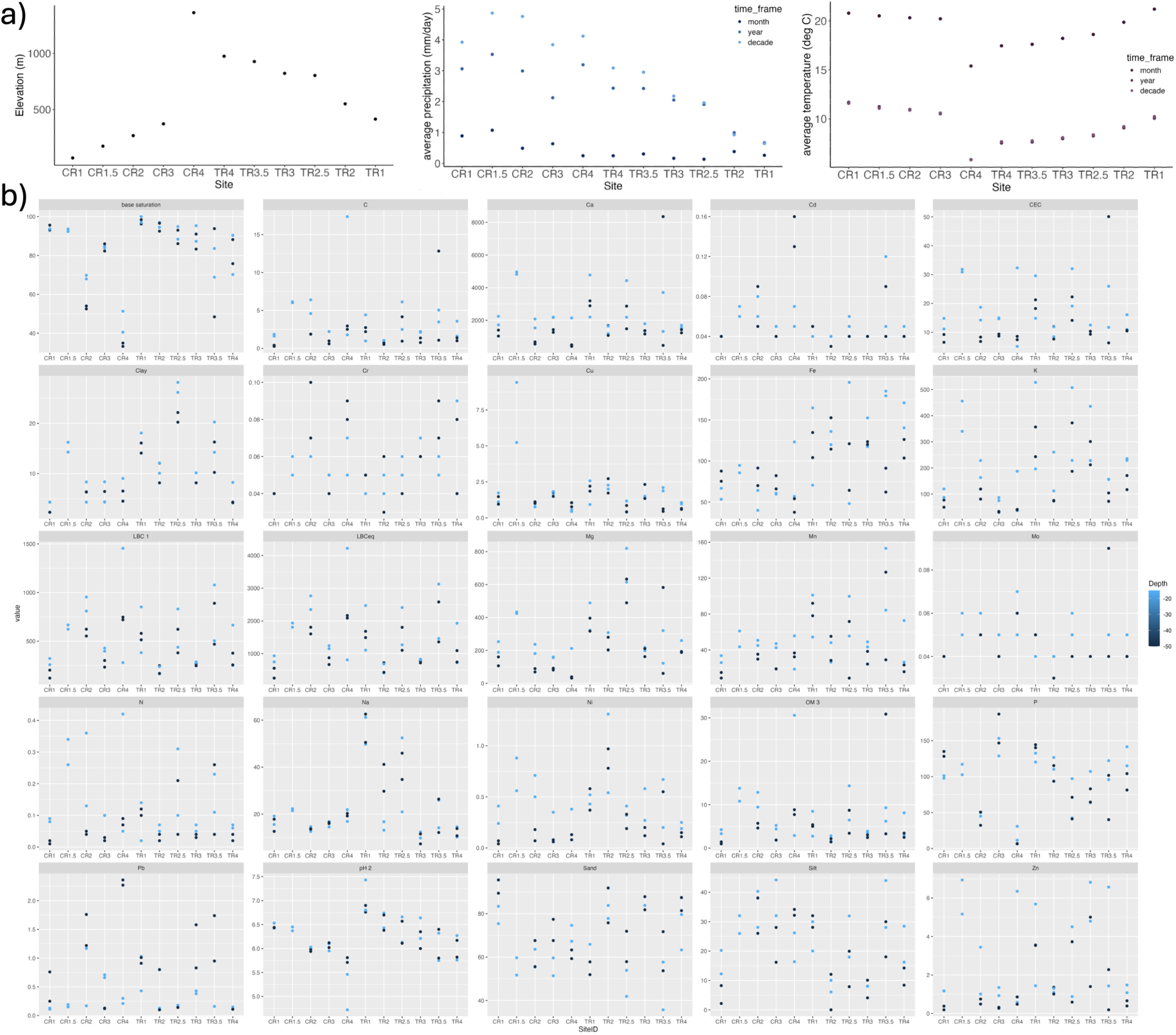
Climate variables by site (a) Site elevation (left), average precipitation for the month, year, and decade leading up to the sampling as predicted by DAYMET (middle), and the average daily temperature in the month, year, and decade leading up to sampling (right). Soil test results (b) colored by depth.

**S3:** The chosen sites represented a wide range of climate conditions, driven primarily by elevation variation, which was inversely correlated with temperature, and precipitation, which was higher on the western side of the mountain range and lower in the east. All sites along the lower Cowlitz River on the western side of the Cascade Mountains had lower elevations than any of the sites on the Tieton River to the east. The highest elevation site was CR4, located at the crest of the White Pass at 1,362 meters. Elevation gradually decreased on the Tieton River side and sharply decreased on the Cowlitz River side to the lowest site at 70 meters, covering a range of ∼1,300 meters. The precipitation patterns varied 10-fold and generally reduced from west to east, with precipitation in the month, year, and decade leading up to sampling being highest at site CR1.5, the second farthest to the west, with an average daily precipitation rate of 4.87 mm/day over the decade, and gradually decreasing to the lowest average precipitation at TR1, the easternmost site, with 0.46 mm/day over the decade. Average temperatures were inversely correlated with the elevation, with a sharp dip at the highest site, CR4. The average daily maximum temperature over the decade ranged from 10.18 C at site CR4 to 17.08 C at site TR1.

Some soil factors had considerable site stochasticity while others correlated with the elevation gradient, primarily those related to biological processes, like carbon to nitrogen ratios, which increase at higher elevations due to reduced decomposition rates, and soil properties affected by leaching, like pH. pH, magnesium content, and copper content were all inversely correlated with elevation. The watershed origin also affected the soil. For example, soil iron content was higher in the Tieton River sites than in the Cowlitz River sites and sodium increased in the two driest, lowest elevation Tieton River sites. The general soil structure also varied across the transect with the Cowlitz River watershed being primarily silt, whereas the Tieton River watershed was dominated by higher sand and clay.

**S4**: Primary PCR reactions contained 12.5 μL of 2× KAPPA HiFi HotStart Ready Mix, 5 μM each of amplicon-specific primers, 25-50 ng of template DNA, and water to 25 μL final volume. For root material, reactions additionally included 0.31 μL each of host peptide nucleic acid blockers at 100 μM (Cregger et al. 2018) purchased from PNA Bio (Thousand Oaks, CA). Thermal cycler conditions for soil samples consisted of initial denaturation at 95°C for 3 min, followed by 25 cycles of 95°C for 30 s, 55°C for 30 s, 72°C for 30 s, and final extension at 72°C for 5 min. Root samples with PNA blockers used 30 cycles with an additional 78°C for 10 s step after the 95°C denaturation.

Following primary amplification, 2 μL of PCR product was verified by gel electrophoresis (1% agarose). The remaining 23 μL was cleaned with AMPure beads (Beckman Coulter, Brea, CA) at 0.8× ratio and eluted in 50 μL of 10 mM Tris pH 8.5. Secondary PCR (Index PCR) reactions contained 25 μL of 2× KAPPA HiFi HotStart Ready Mix, 5 μL each of Nextera XT Index primers (N7xx and S5xx), 5 μL of purified primary PCR product, and 10 μL water in 50 μL total volume. Thermal cycler conditions consisted of denaturation at 95°C for 3 min, followed by 8 cycles of 95°C for 30 s, 55°C for 30 s, 72°C for 30 s, and final extension at 72°C for 5 min. Indexed amplicons were cleaned with AMPure beads (0.8x ratio) and resuspended in 25 μL of 10 mM Tris pH 8.5. Products were quantified on a Nanodrop instrument (ThermoFisher, Waltham, MA), pooled equimolarly, and validated on an Agilent Bioanalyzer (Agilent, Santa Clara, CA) using a DNA7500 chip. The pool was purified again with AMPure beads (0.8x ratio) to remove small DNA fragments, and final concentration was determined using a Qubit instrument (Life Technologies, Carlsbad, CA) with the broad range double-stranded DNA assay. Libraries were denatured with 0.2 N sodium hydroxide, diluted to final sequencing concentration, and spiked with PhiX control DNA.

**S5:** RNA was extracted from foliar samples by adding 100 mg of ground, cryo-frozen foliar tissue to 850 uL CTAB buffer and 10 uL B-mercaptoethanol, vortexed for 2 minutes, and incubated at 30C while shaking at 1800 rpm 5 minutes, then digested in 600 uL of 24:1 chloroform-isoamylalcohol. Samples were vortexed for 1 minute and centrifuged for 8 minutes at maximum speed. The top layer of supernatant (about 780 uLs) was pipetted from the sample and applied to the appropriate well of a Promega Maxwell® RSC Plant RNA kit cartridge. The subsequent RNA purification, DNase treatment and RNA elution steps were done on a Maxwell® RSC Instrument according to the kit protocol.

Complementary DNA (cDNA) was synthesized from total RNA for qRT-PCR using igScript™ (IG Intact Genomics). Each reaction included 1ug total RNA, 4ul 5x igScript master mix, 2ul 60uM random hexamer primer and sterile water to 20ul. Synthesis reactions were carried out on a thermocycler at 25C for 10 min, 42C for 60 min, 65C for 20 min and held at 4C until removal. qRT-PCR was performed with the following cycling conditions: initial denaturation at 95C for 10 minutes, 40 amplification cycles consisting of 95C denature for 30 seconds, 60C annealing for 1 minute, followed by a 50 to 95C melt curve. Samples were run in triplicate in 10 uL reactions consisting of 5 uL SYBR Green PCR Master Mix (ThermoFisher), 0.5 uM F and R primers, and 1 uM of 1:10 diluted cDNA.

**S6:**
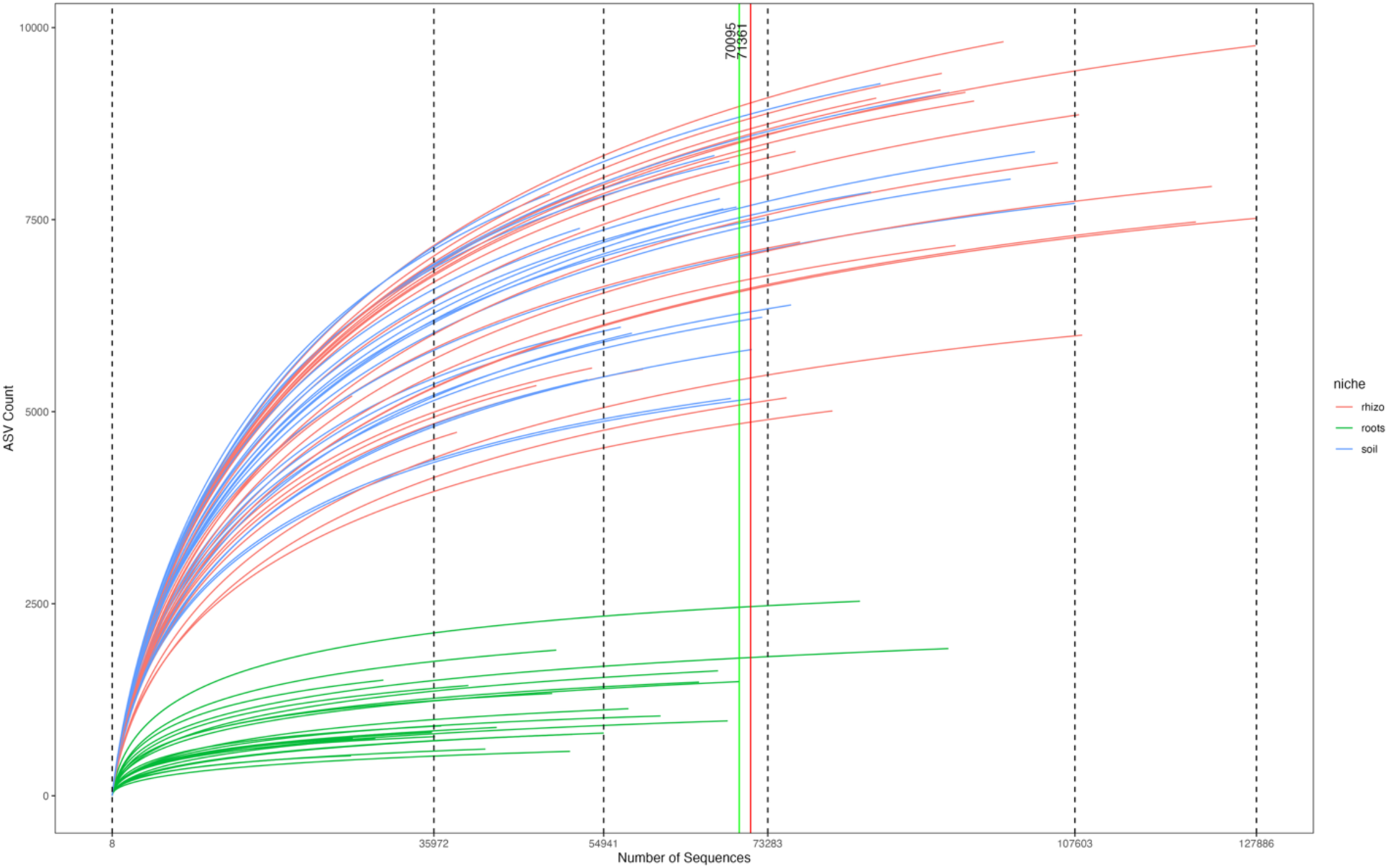
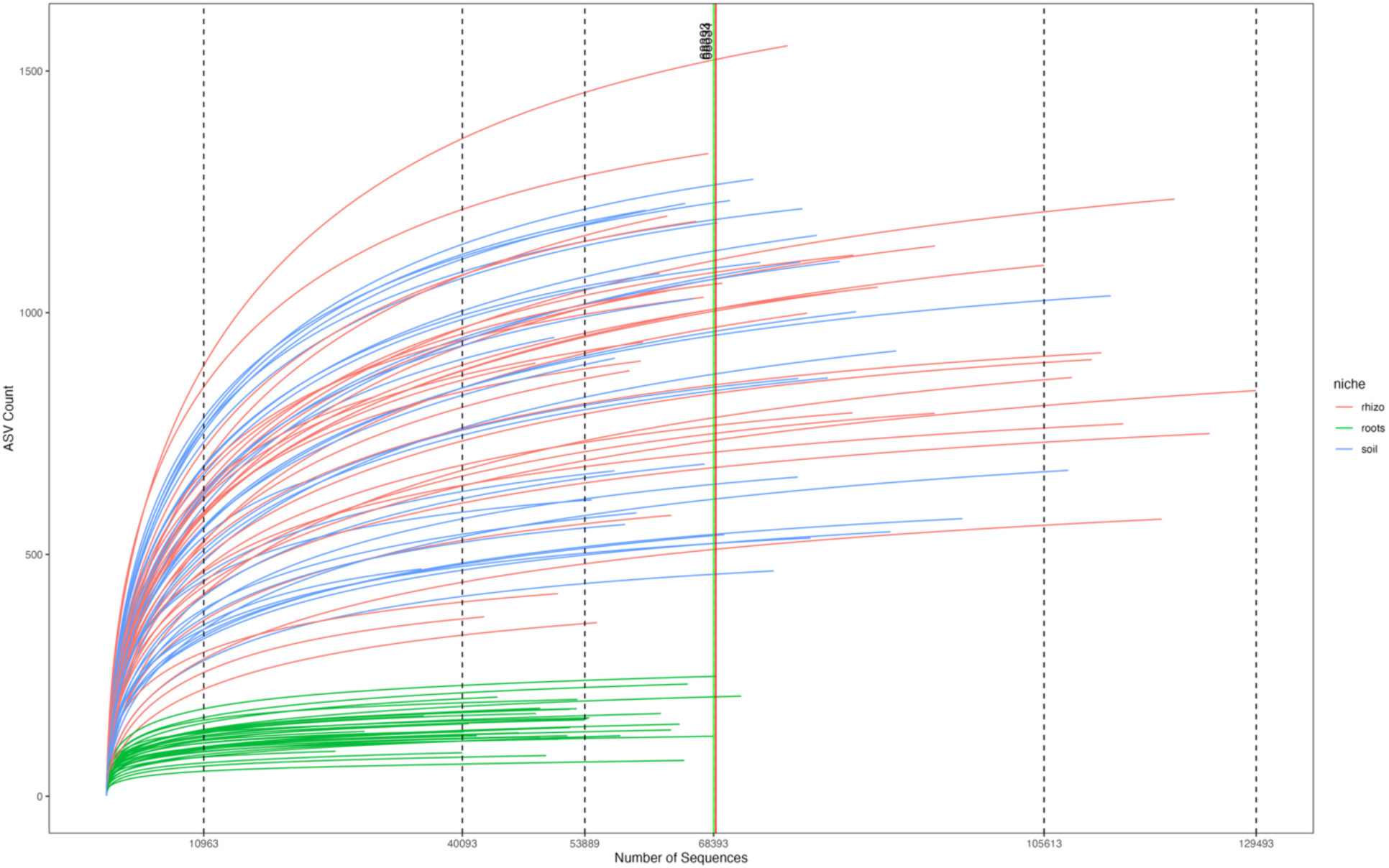
Rarefaction curve plots for the 16S library (top) and ITS2 (bottom). The colors indicate the sample type with bulk soil shown in blue, rhizosphere soil in red, and root endospheres in green. The dotted vertical lines are at 10%, 30%, 60%, 90%, and 100% of the unrarefied sequence counts. The green line represents the cutoff for a minimum replication of three samples per niche compartment and the red line indicates the cutoff for a minimum replication of two samples.

**S7:**
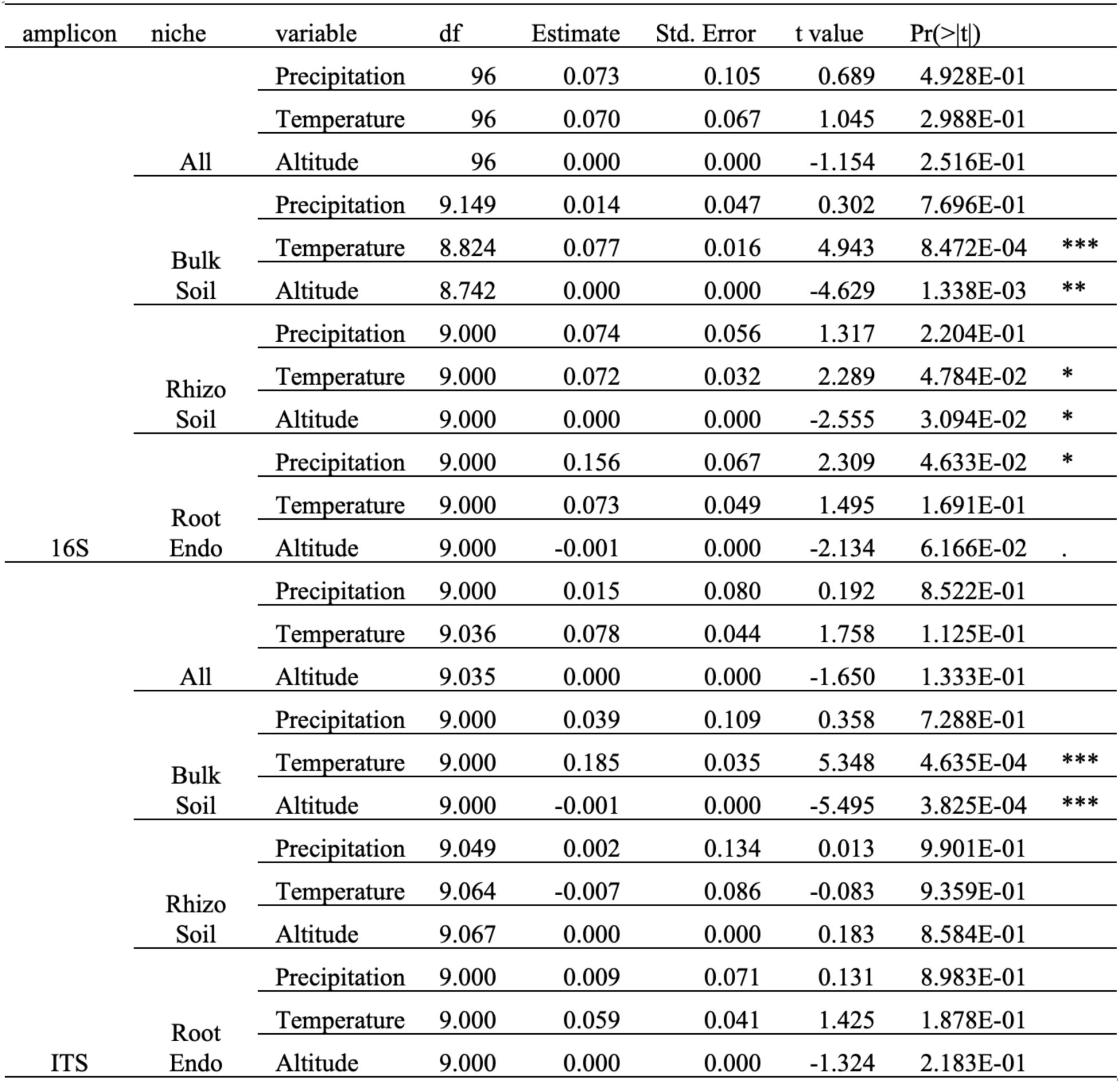
Linear Mixed Effects Models (LMEs) showing the correlation between continuous climate variables extracted from DAYMET based on site coordinates and bacterial/archaeal and fungal Shannon Diversity Index with site as a random factor to control for pseudoreplication. Asterisks indicate statistically significant correlations.

**S8:**
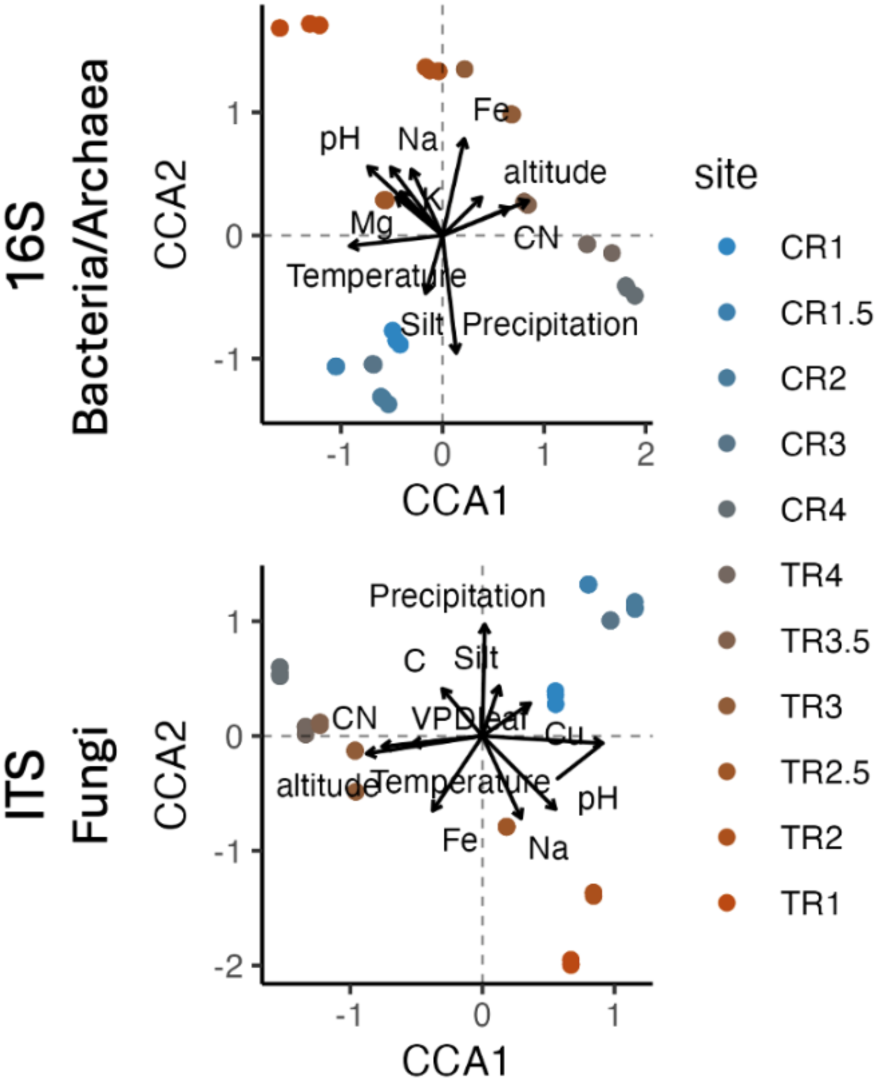
Canonical Correspondence Analysis (CCA) plots for 16S/Bacteria/Archaea (top) and ITS/Fungi (bottom) for the full sample set, showing the influence of climate (average rainfall and daily maximum temperature in the decade leading up to sampling), altitude, soil factors (pH, Fe, CN, Na, Clay, C, Silt, Mg, Cu, K, OM, P), and *Populus* host factors (VPDleaf, Tleaf, gsw). Points are color-coded by site from the Cowlitz River to the Tieton river on a blue to red scale.

**S9:**
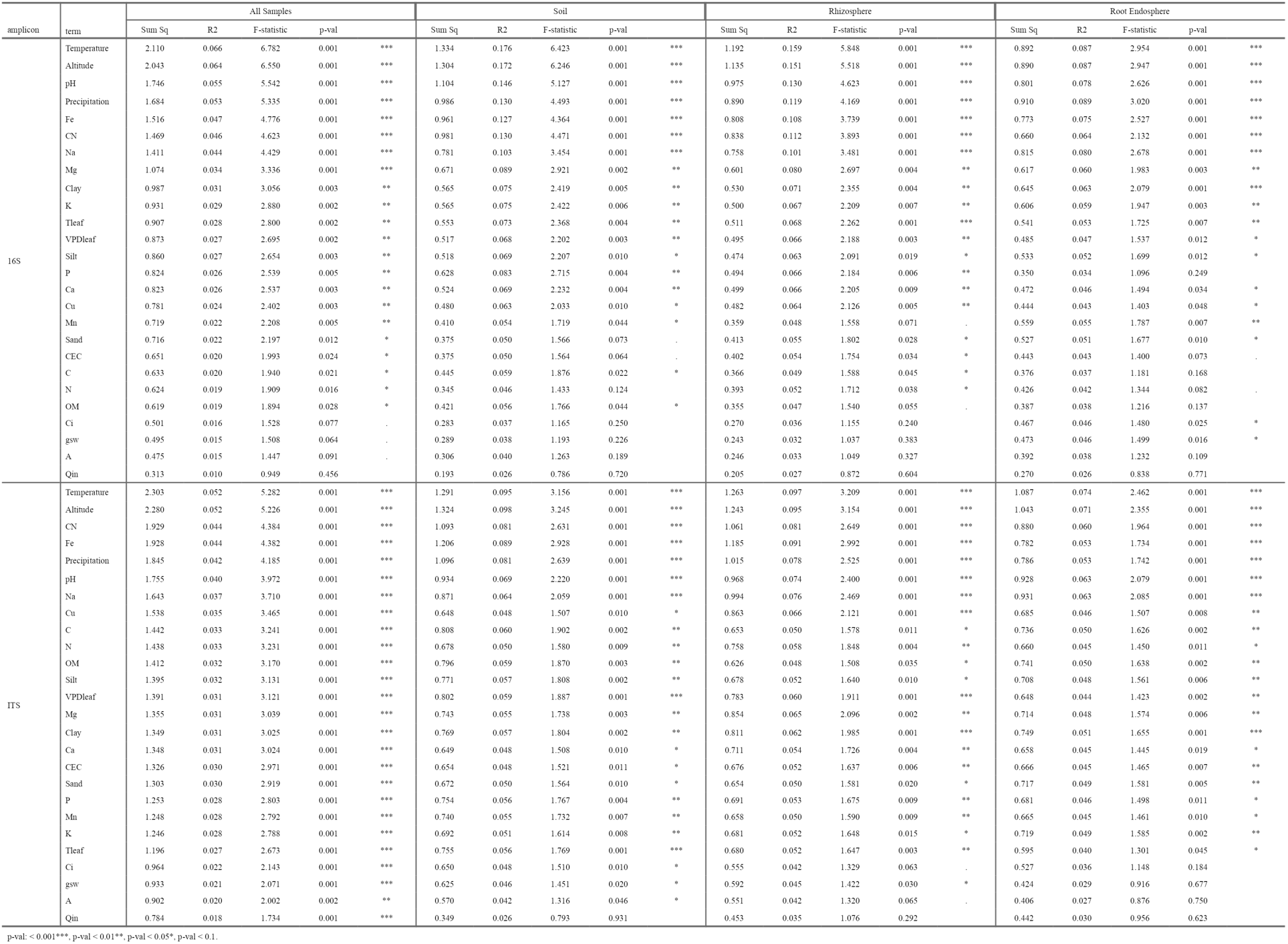
Adonis rank correlations testing the correlation of each climate, soil, and tree phenotype metric and the microbial beta-diversity. Ranks were used to inform linear mixed-effects (LME) model construction.

**S10:**
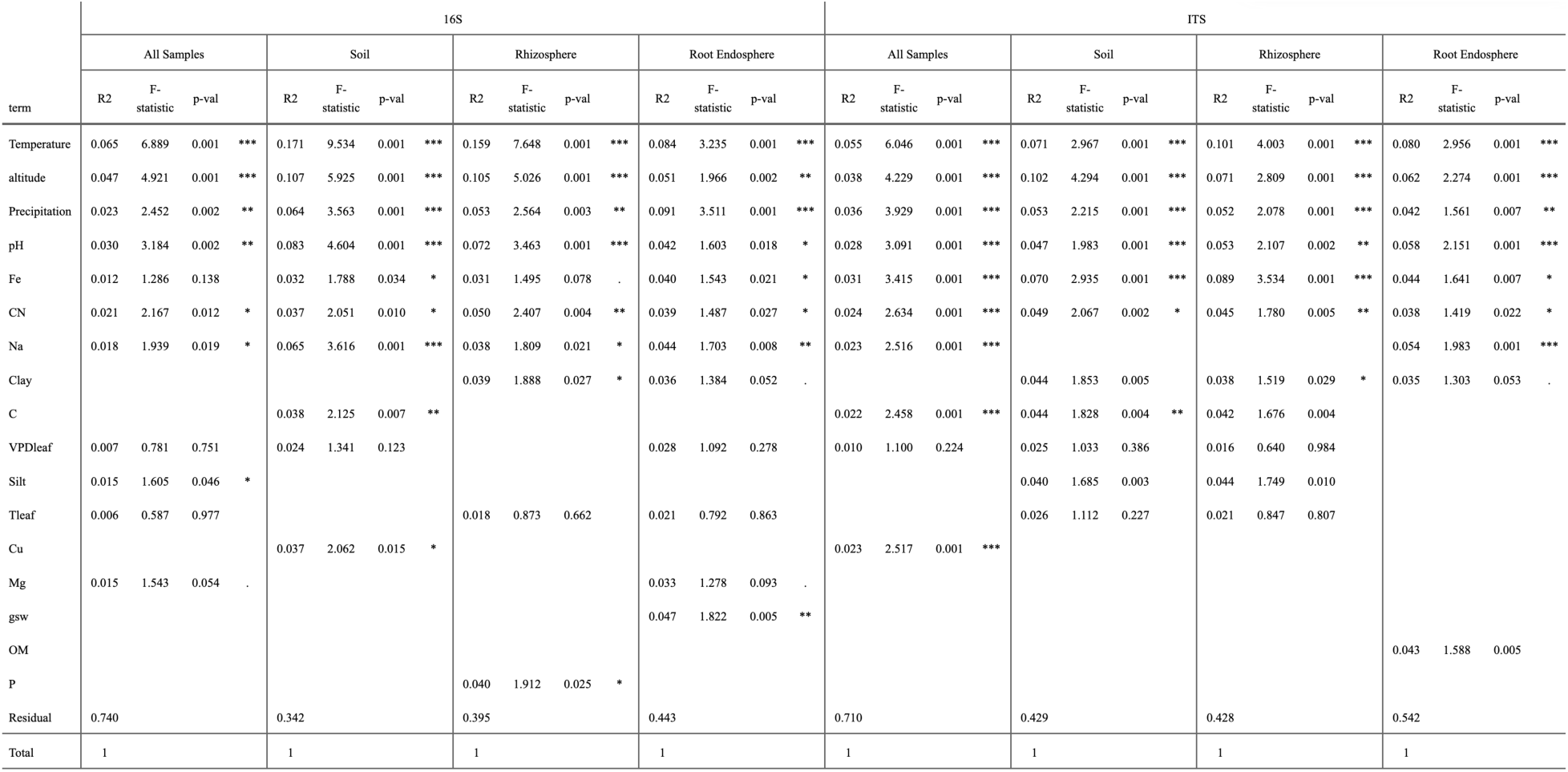
Linear mixed-effects models (LMEs) of the effects of climate (average rainfall and temperature in the decade leading up to sampling), altitude, soil factors (pH, Fe, CN, Na, Clay, C, Silt, Mg, Cu, K, OM, P), and *P. trichocarpa* host factors (VPDleaf, Tleaf, gsw) on the bacterial/archaeal and fungal beta-diversity in all samples, bulk soil, rhizosphere soil, and root endosphere. Site CR1.5 was excluded as only shallow cores were taken due to rocky conditions.

**S11:**
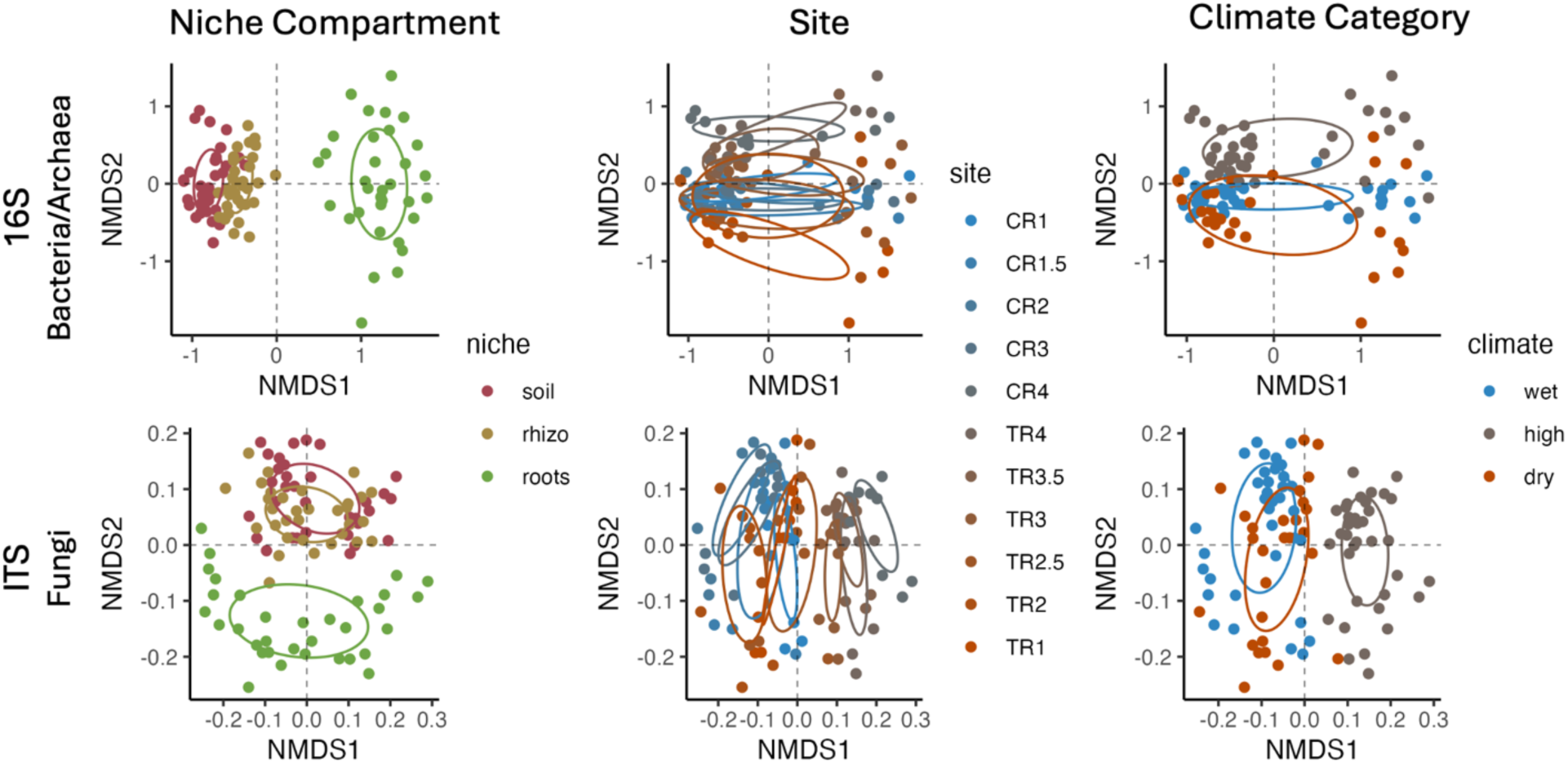
NMDS plots showing the beta-diversity of all bacterial/archaeal samples (top) and all fungal samples (bottom). Plots are color-coded by niche compartment (left) with bulk soil in red, rhizosphere soil in yellow, and root endosphere in green, site (center) with a color gradient from blue in the wetter sites to red in the drier sites across the transect, and climate category (right) with blue representing the cluster of wetter sites, grey representing the cluster of colder, high-altitude sites, and red representing the cluster of dry sites.

**S12:**
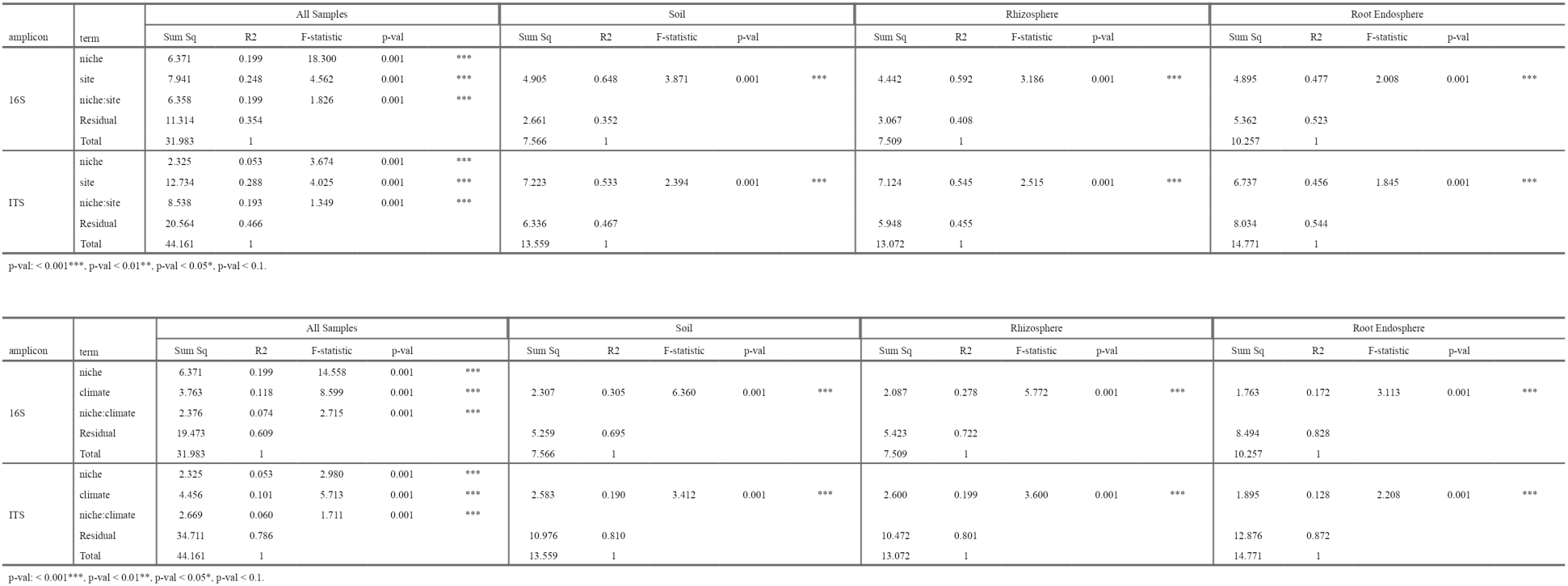
Top: PERMANOVA of niche and site on 16S-bacterial/archaeal and ITS-fungal beta-diversity for niche, site, and niche:site for all samples, bulk soil, rhizosphere soil, and the root endosphere compartment. Bottom: PERMANOVA tables showing the impact of niche and climate category (wet, high, dry) on bacterial/archaeal and fungal beta-diversity.

| amplicon | term | All Samples |  |  |  | Soil |  |  |  | Rhizosphere |  |  |  | Root Endosphere |  |  |  |  |  |  |  |
| --- | --- | --- | --- | --- | --- | --- | --- | --- | --- | --- | --- | --- | --- | --- | --- | --- | --- | --- | --- | --- | --- |
|  |  | Sum Sq | R2 | F-statistic | p-val | Sum Sq | R2 | F-statistic | p-val | Sum Sq | R2 | F-statistic | p-val | Sum Sq | R2 | F-statistic | p-val |  |  |  |  |
| 16S | niche | 6.371 | 0.199 | 18.300 | 0.001 | *** |  |  |  |  |  |  |  |  |  |  |  |  |  |  |  |
|  | site | 7.941 | 0.248 | 4.562 | 0.001 | *** | 4.905 | 0.648 | 3.871 | 0.001 | *** | 4.442 | 0.592 | 3.186 | 0.001 | *** | 4.895 | 0.477 | 2.008 | 0.001 | *** |
|  | niche:site | 6.358 | 0.199 | 1.826 | 0.001 | *** |  |  |  |  |  |  |  |  |  |  |  |  |  |  |  |
|  | Residual | 11.314 | 0.354 |  |  |  | 2.661 | 0.352 |  |  |  | 3.067 | 0.408 |  |  |  | 5.362 | 0.523 |  |  |  |
|  | Total | 31.983 | 1 |  |  |  | 7.566 | 1 |  |  |  | 7.509 | 1 |  |  |  | 10.257 | 1 |  |  |  |
| ITS | niche | 2.325 | 0.053 | 3.674 | 0.001 | *** |  |  |  |  |  |  |  |  |  |  |  |  |  |  |  |
|  | site | 12.734 | 0.288 | 4.025 | 0.001 | *** | 7.223 | 0.533 | 2.394 | 0.001 | *** | 7.124 | 0.545 | 2.515 | 0.001 | *** | 6.737 | 0.456 | 1.845 | 0.001 | *** |
|  | niche:site | 8.538 | 0.193 | 1.349 | 0.001 | *** |  |  |  |  |  |  |  |  |  |  |  |  |  |  |  |
|  | Residual | 20.564 | 0.466 |  |  |  | 6.336 | 0.467 |  |  |  | 5.948 | 0.455 |  |  |  | 8.034 | 0.544 |  |  |  |
|  | Total | 44.161 | 1 |  |  |  | 13.559 | 1 |  |  |  | 13.072 | 1 |  |  |  | 14.771 | 1 |  |  |  |

p-val: < 0.001\*\*\*, p-val < 0.01\*\*, p-val < 0.05\*, p-val < 0.1. p-val < 0.001\*\*\*, p-val < 0.01\*\*, p-val < 0.05\*, p-val < 0.1.
| amplicon | term | All Samples |  |  |  | Soil |  |  |  | Rhizosphere |  |  |  | Root Endosphere |  |  |  |  |  |  |  |
| --- | --- | --- | --- | --- | --- | --- | --- | --- | --- | --- | --- | --- | --- | --- | --- | --- | --- | --- | --- | --- | --- |
|  |  | Sum Sq | R2 | F-statistic | p-val | Sum Sq | R2 | F-statistic | p-val | Sum Sq | R2 | F-statistic | p-val | Sum Sq | R2 | F-statistic | p-val |  |  |  |  |
| 16S | niche | 6.371 | 0.199 | 14.558 | 0.001 | *** |  |  |  |  |  |  |  |  |  |  |  |  |  |  |  |
|  | climate | 3.763 | 0.118 | 8.599 | 0.001 | *** | 2.307 | 0.305 | 6.360 | 0.001 | *** | 2.087 | 0.278 | 5.772 | 0.001 | *** | 1.763 | 0.172 | 3.113 | 0.001 | *** |
|  | niche:climate | 2.376 | 0.074 | 2.715 | 0.001 | *** |  |  |  |  |  |  |  |  |  |  |  |  |  |  |  |
|  | Residual | 19.473 | 0.609 |  |  |  | 5.259 | 0.695 |  |  |  | 5.423 | 0.722 |  |  |  | 8.494 | 0.828 |  |  |  |
|  | Total | 31.983 | 1 |  |  |  | 7.566 | 1 |  |  |  | 7.509 | 1 |  |  |  | 10.257 | 1 |  |  |  |
| ITS | niche | 2.325 | 0.053 | 2.980 | 0.001 | *** |  |  |  |  |  |  |  |  |  |  |  |  |  |  |  |
|  | climate | 4.456 | 0.101 | 5.713 | 0.001 | *** | 2.583 | 0.190 | 3.412 | 0.001 | *** | 2.600 | 0.199 | 3.600 | 0.001 | *** | 1.895 | 0.128 | 2.208 | 0.001 | *** |
|  | niche:climate | 2.669 | 0.060 | 1.711 | 0.001 | *** |  |  |  |  |  |  |  |  |  |  |  |  |  |  |  |
|  | Residual | 34.711 | 0.786 |  |  |  | 10.976 | 0.810 |  |  |  | 10.472 | 0.801 |  |  |  | 12.876 | 0.872 |  |  |  |
|  | Total | 44.161 | 1 |  |  |  | 13.559 | 1 |  |  |  | 13.072 | 1 |  |  |  | 14.771 | 1 |  |  |  |

p-val < 0.001\*\*\*, p-val < 0.01\*\*, p-val < 0.05\*, p-val < 0.1.

**S13:**
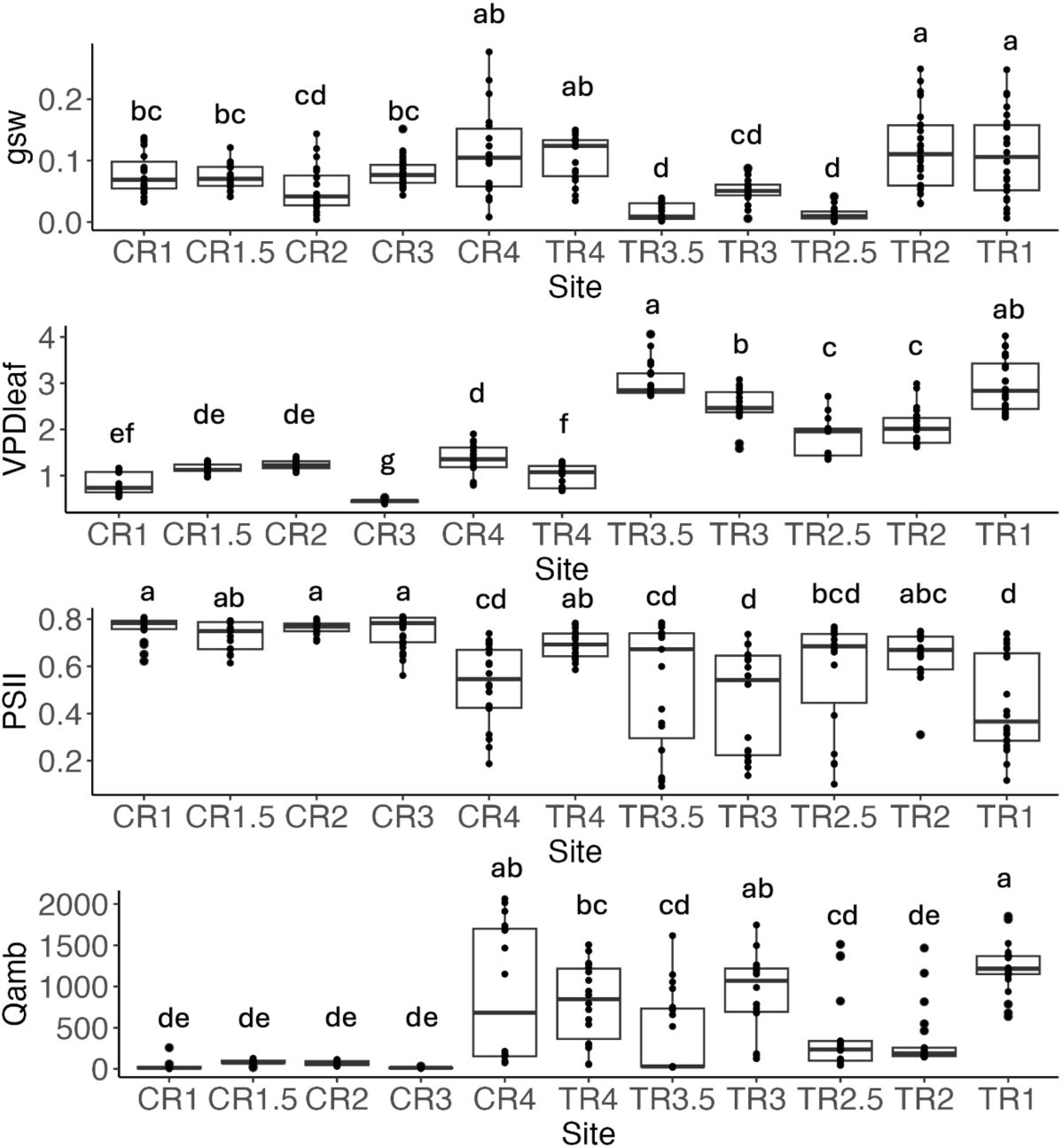
Tree phenotype data for (top to bottom) stomatal conductance to water vapor in mol m^-2^ s^-1^ (gsw), vapor pressure deficit at the temperature of the leaf in kPa (VPDleaf), photosystem II in 1 – steady state flux / maximum flux (PSII), and ambient light in µmol of photons m⁻² s⁻¹ (Qamb) by site. Letters indicate Tukey HSD groups.

**S14:**
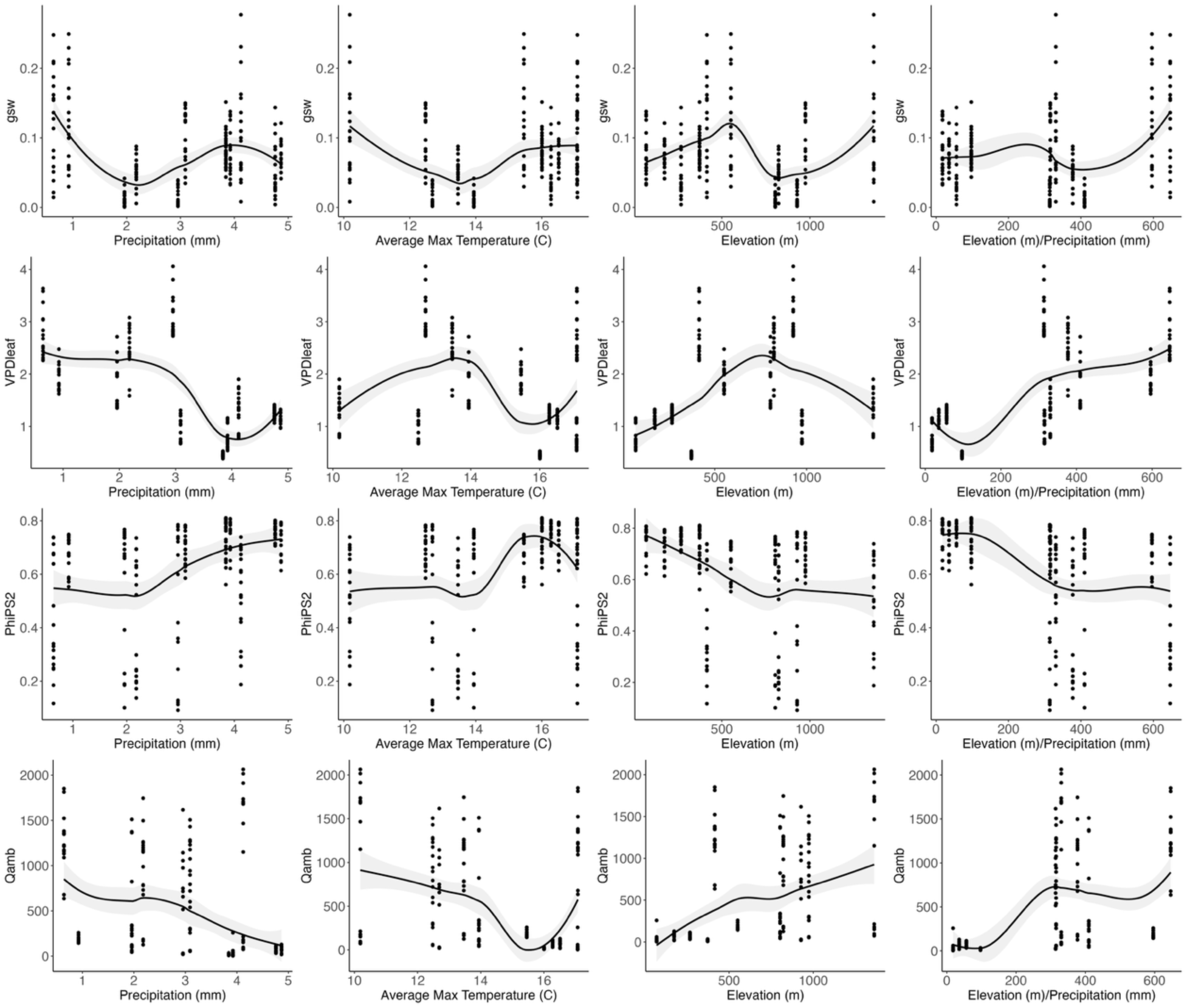
Tree photosynthesis data correlated with environmental variables calculated in DAYMET.

**S15:** Table of the Pearson Correlations between plant phenotype metrics as determined by LICOR and climate metrics collected from DAYMET.

|  | r | p-val |  |
| --- | --- | --- | --- |
| gsw by Precipitation | -0.139 | 0.043 | * |
| gsw by Temperature | 0.070 | 0.311 |  |
| gsw by Elevation | -0.019 | 0.782 |  |
| VPDleaf by Precipitation | -0.612 | 0.000 | *** |
| VPDleaf by Temperature | -0.212 | 0.002 | ** |
| VPDleaf by Elevation | 0.329 | 0.000 | *** |
| PhiPS2 by Precipitation | 0.371 | 0.000 | *** |
| PhiPS2 by Temperature | 0.297 | 0.000 | *** |
| PhiPS2 by Elevation | -0.379 | 0.000 | *** |
| Qamb by Precipitation | -0.389 | 0.000 | *** |
| Qamb by Temperature | -0.352 | 0.000 | *** |
| Qamb by Elevation | 0.453 | 0.000 | *** |
| gsw by Elevation/Precipitation | 0.016 | 0.821 |  |
| VPDleaf by Elevation/Precipitation | 0.245 | 0.000 | *** |
| PhiPS2 by Elevation/Precipitation | -0.345 | 0.000 | *** |
| Qamb by Elevation/Precipitation | 0.430 | 0.000 | *** |

p-val: < 0.001\*\*\*, p-val < 0.01\*\*, p-val < 0.05\*, p-val < 0.1.

**S16:**
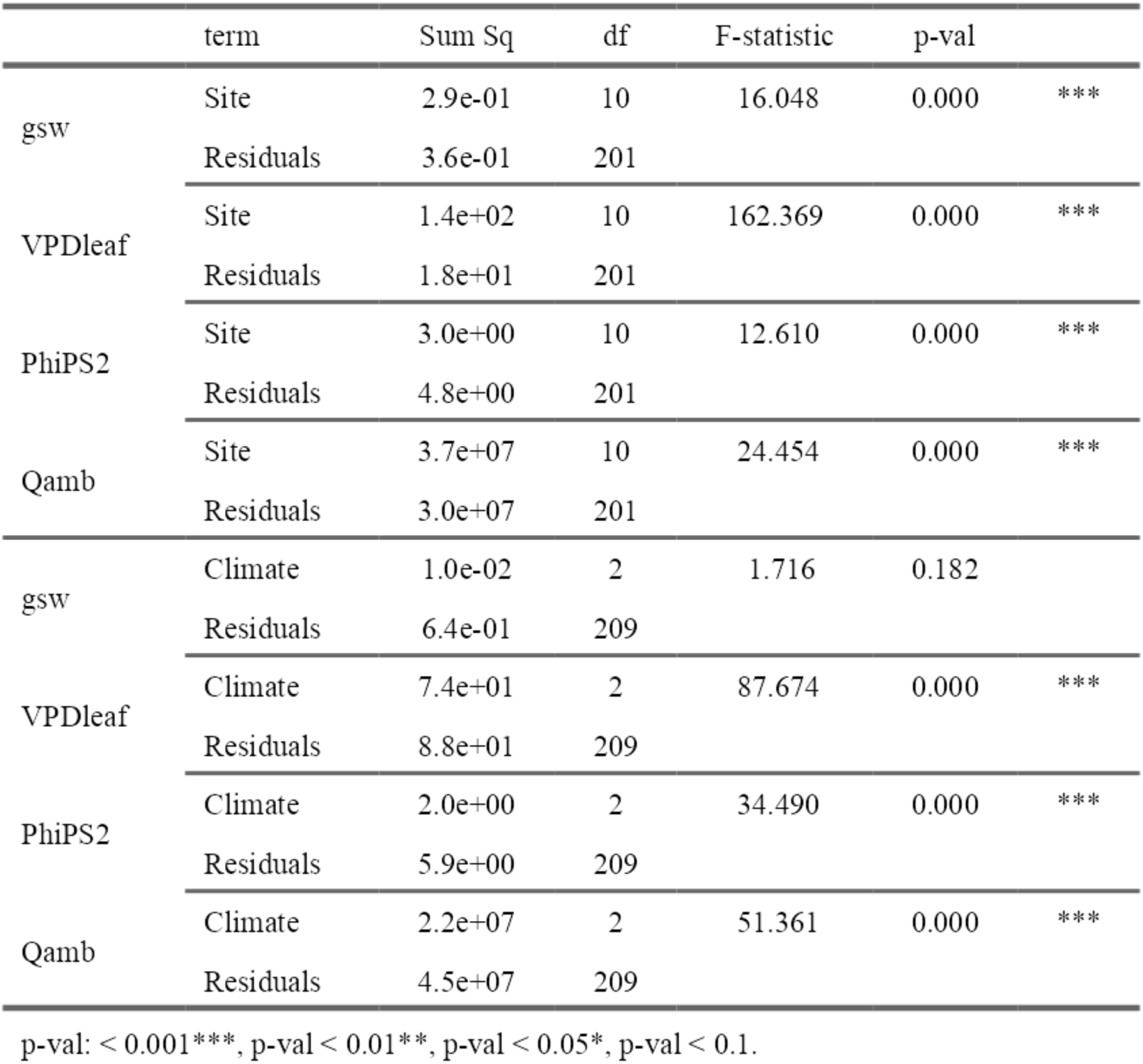
Table displaying ANOVAs evaluating correlations between site and plant photosynthesis and climate and plant photosynthesis.

|  | term | Sum Sq | df | F-statistic | p-val |  |
| --- | --- | --- | --- | --- | --- | --- |
| gsw | Site | 2.9e-01 | 10 | 16.048 | 0.000 | *** |
|  | Residuals | 3.6e-01 | 201 |  |  |  |
| VPDleaf | Site | 1.4e+02 | 10 | 162.369 | 0.000 | *** |
|  | Residuals | 1.8e+01 | 201 |  |  |  |
| PhiPS2 | Site | 3.0e+00 | 10 | 12.610 | 0.000 | *** |
|  | Residuals | 4.8e+00 | 201 |  |  |  |
| Qamb | Site | 3.7e+07 | 10 | 24.454 | 0.000 | *** |
|  | Residuals | 3.0e+07 | 201 |  |  |  |
| gsw | Climate | 1.0e-02 | 2 | 1.716 | 0.182 |  |
|  | Residuals | 6.4e-01 | 209 |  |  |  |
| VPDleaf | Climate | 7.4e+01 | 2 | 87.674 | 0.000 | *** |
|  | Residuals | 8.8e+01 | 209 |  |  |  |
| PhiPS2 | Climate | 2.0e+00 | 2 | 34.490 | 0.000 | *** |
|  | Residuals | 5.9e+00 | 209 |  |  |  |
| Qamb | Climate | 2.2e+07 | 2 | 51.361 | 0.000 | *** |
|  | Residuals | 4.5e+07 | 209 |  |  |  |

p-val: < 0.001\*\*\*, p-val < 0.01\*\*, p-val < 0.05\*, p-val < 0.1.

**S17:**
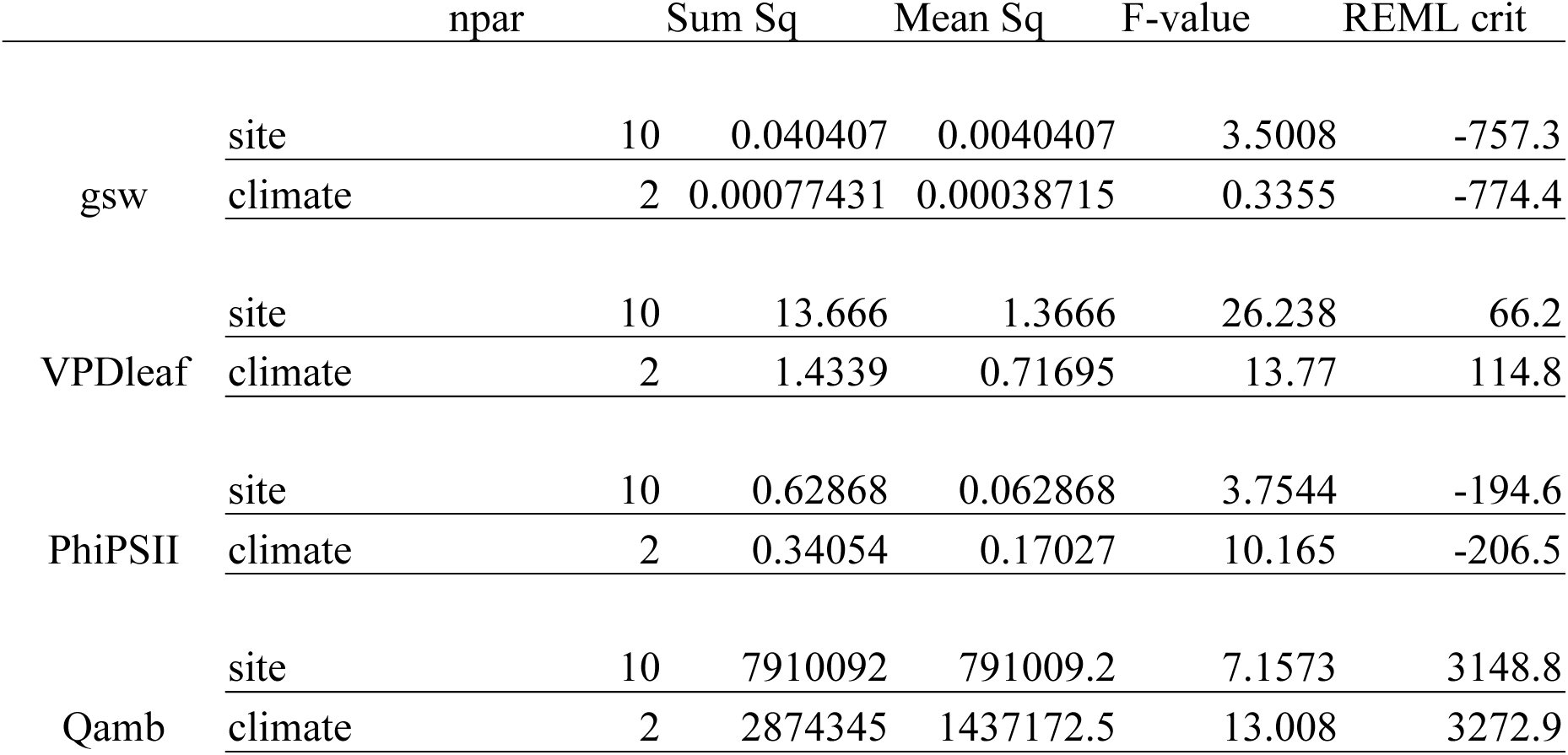
ANOVA tables of linear mixed effects models, to control for within tree sampling effects. Site was significantly correlated with gsw, VPDleaf, PhiPSII, and Qamb. Climate was significantly correlated with VPDleaf, and was more strongly correlated with PhiPSII and Qamb than site when controlling for the tree sampling effect.

**S18:**
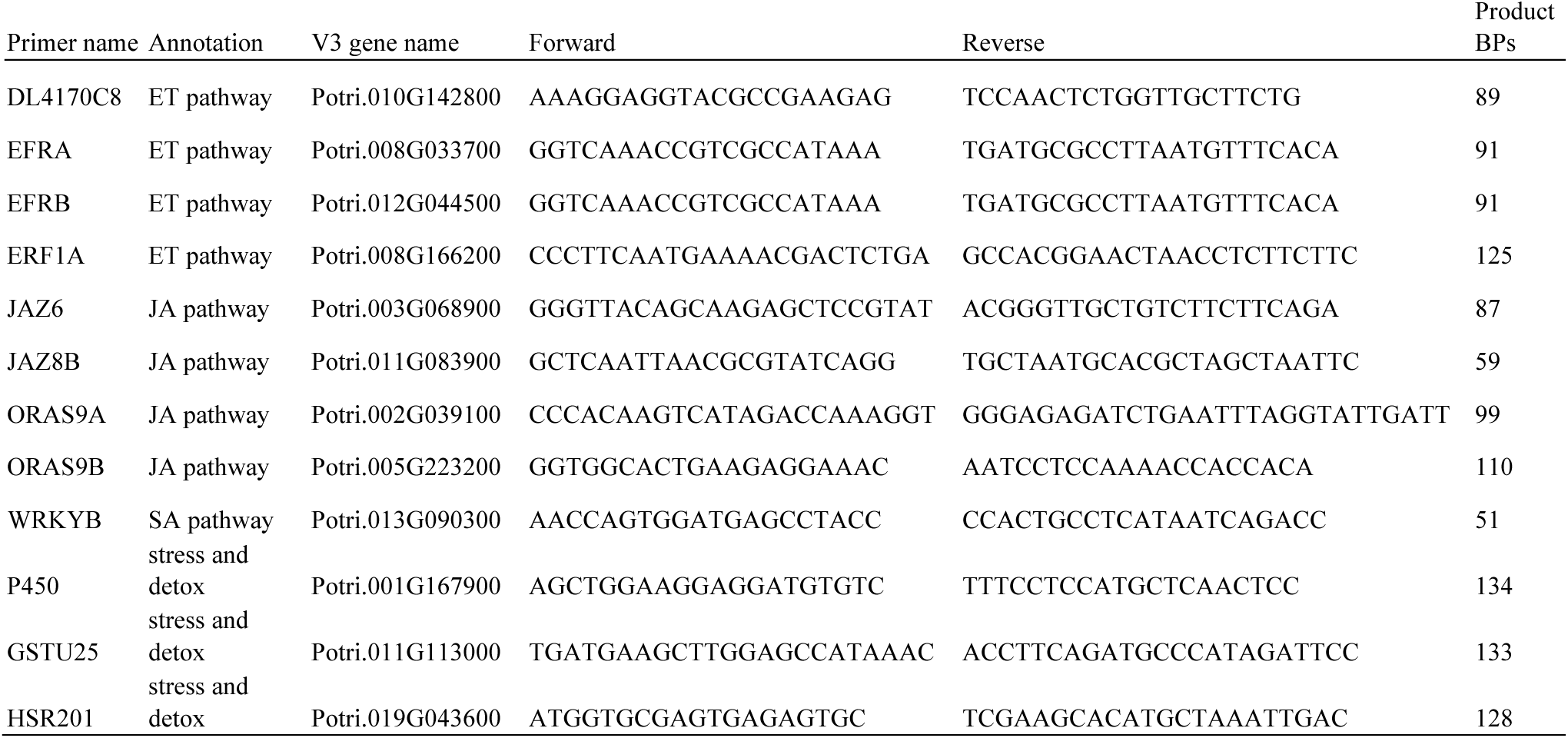
Table with information about the foliar abiotic stress expression primers and target genes for chosen for this study.

**S19:**
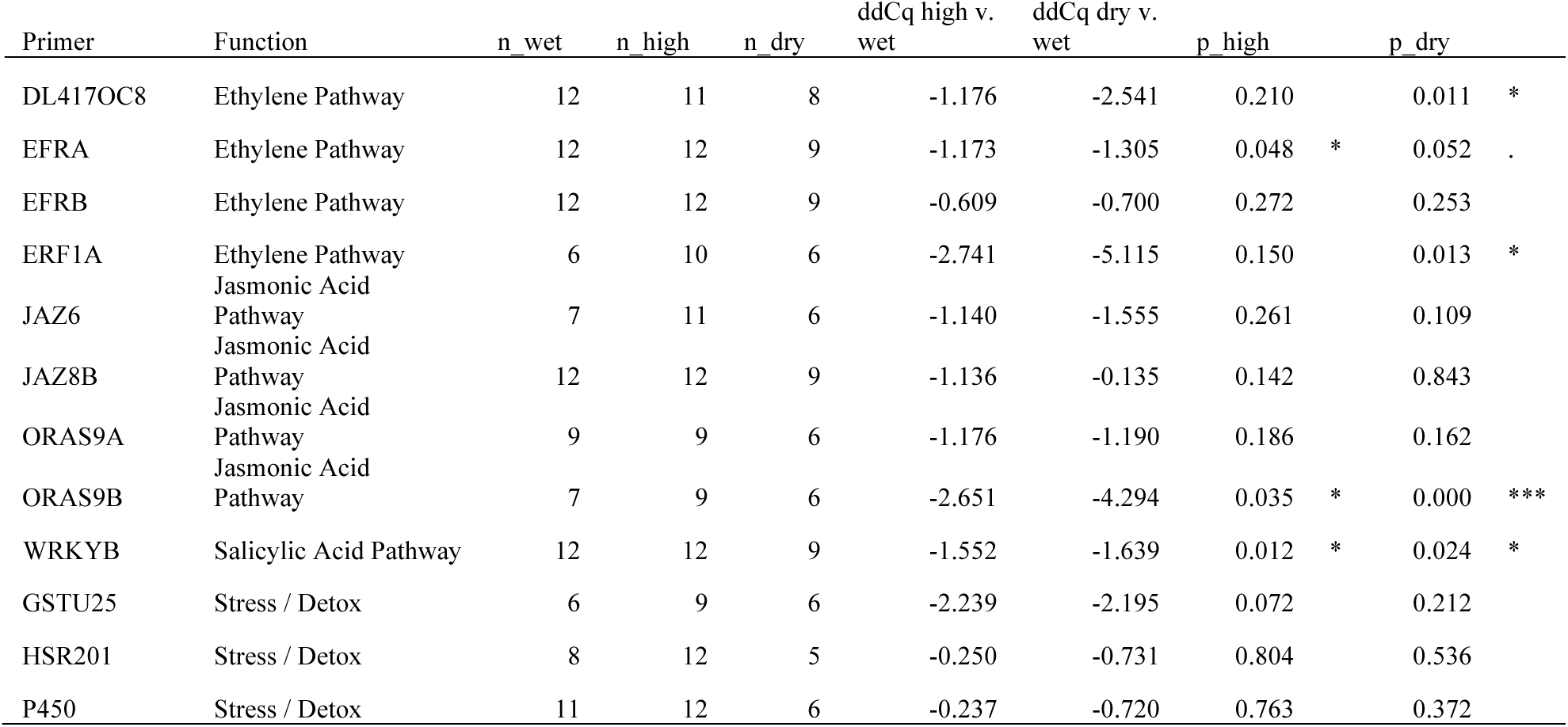
Table of the qRT-PCR results. ddCq values were calculated against housekeeping gene expression (UBQ10B and 18S). n columns indicate the number of samples from each climate treatment that successfully replicated out of 12 total wet samples, 12 total high samples, and 9 total dry samples. The average expression across the wet sites was used as a control. Negative ddCqs indicate higher expression in the high or dry sites compared to the wet sites. P-values were determined with student t-tests.

**S20:**
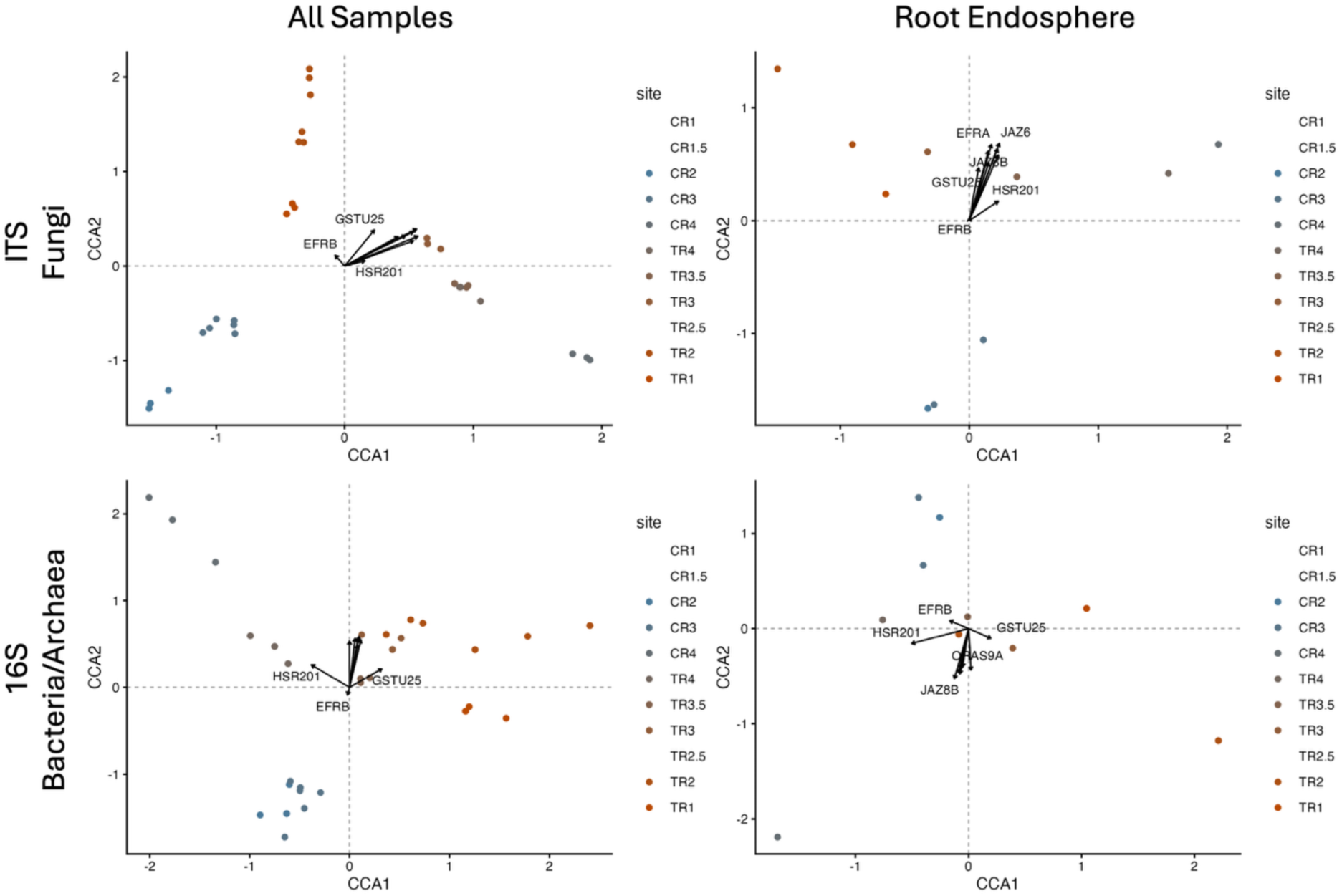
CCA plots showing correlations between genes and beta-diversity of the fungal communities (top) and the bacterial/archaeal communities (bottom) of the full sample set (left) and the root endosphere (right).

**S21:**
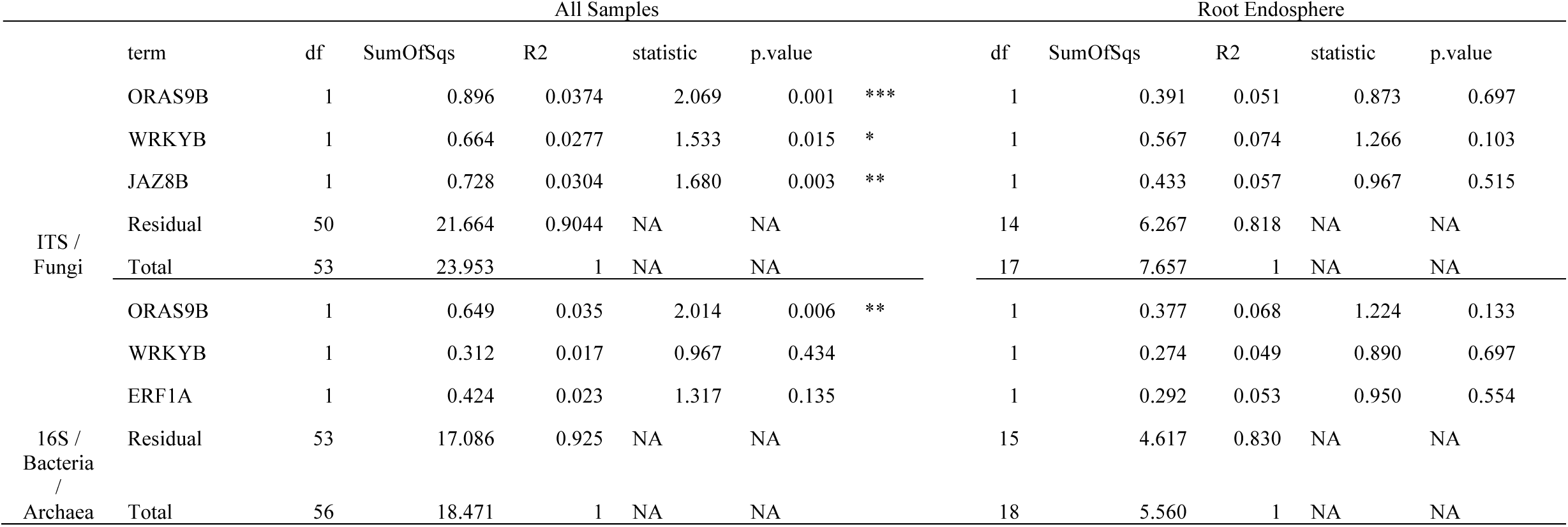
Table showing PERMANOVAs of the qRT-PCR results. Model construction was informed with individual gene vs. microbial beta-diversity pearson correlations.

| All Samples |  |  |  |  |  |  | Root Endosphere |  |  |  |  |  |
| --- | --- | --- | --- | --- | --- | --- | --- | --- | --- | --- | --- | --- |
|  | term | df | SumOfSqs | R2 | statistic | p.value |  | df | SumOfSqs | R2 | statistic | p.value |
| ITS /<br>Fungi | ORAS9B | 1 | 0.896 | 0.0374 | 2.069 | 0.001 *** |  | 1 | 0.391 | 0.051 | 0.873 | 0.697 |
|  | WRKYB | 1 | 0.664 | 0.0277 | 1.533 | 0.015 * |  | 1 | 0.567 | 0.074 | 1.266 | 0.103 |
|  | JAZ8B | 1 | 0.728 | 0.0304 | 1.680 | 0.003 ** |  | 1 | 0.433 | 0.057 | 0.967 | 0.515 |
|  | Residual | 50 | 21.664 | 0.9044 | NA | NA |  | 14 | 6.267 | 0.818 | NA | NA |
|  | Total | 53 | 23.953 | 1 | NA | NA |  | 17 | 7.657 | 1 | NA | NA |
| 16S /<br>Bacteria<br>/<br>Archaea | ORAS9B | 1 | 0.649 | 0.035 | 2.014 | 0.006 ** |  | 1 | 0.377 | 0.068 | 1.224 | 0.133 |
|  | WRKYB | 1 | 0.312 | 0.017 | 0.967 | 0.434 |  | 1 | 0.274 | 0.049 | 0.890 | 0.697 |
|  | ERF1A | 1 | 0.424 | 0.023 | 1.317 | 0.135 |  | 1 | 0.292 | 0.053 | 0.950 | 0.554 |
|  | Residual | 53 | 17.086 | 0.925 | NA | NA |  | 15 | 4.617 | 0.830 | NA | NA |
|  | Total | 56 | 18.471 | 1 | NA | NA |  | 18 | 5.560 | 1 | NA | NA |

**S22:** Table showing the number of nodes and edges in each network as well as the modularity (the prevalence of distinct, highly connected clusters in relation to the overall connectedness of the network), transitivity (clustering coefficient of the network as a whole), and density (number of edges in the network compared to the total number of possible edges) of each network.

| Climate | Niche | Total Nodes | Bacterial Nodes | Fungal Nodes | Edges | Modularity | Transitivity | Density |
| --- | --- | --- | --- | --- | --- | --- | --- | --- |
| Wet | Bulk Soil | 250 | 96 | 153 | 522 | 0.5819443 | 0.10436137 | 0.01677108 |
|  | Rhizosphere Soil | 250 | 96 | 153 | 537 | 0.5559197 | 0.08312716 | 0.01725301 |
|  | Root Endosphere | 250 | 148 | 102 | 570 | 0.5446645 | 0.10120783 | 0.01831325 |
| High | Bulk Soil | 250 | 95 | 155 | 605 | 0.5345195 | 0.09151047 | 0.01943775 |
|  | Rhizosphere Soil | 250 | 96 | 154 | 535 | 0.5472373 | 0.08697789 | 0.01718876 |
|  | Root Endosphere | 250 | 132 | 118 | 489 | 0.6130055 | 0.16083916 | 0.01571084 |
| Dry | Bulk Soil | 250 | 99 | 147 | 595 | 0.5731827 | 0.12000000 | 0.01911647 |
|  | Rhizosphere Soil | 250 | 123 | 124 | 600 | 0.5545250 | 0.10655091 | 0.01927711 |
|  | Root Endosphere | 250 | 139 | 111 | 570 | 0.5490320 | 0.09087031 | 0.01831325 |

**S23:**
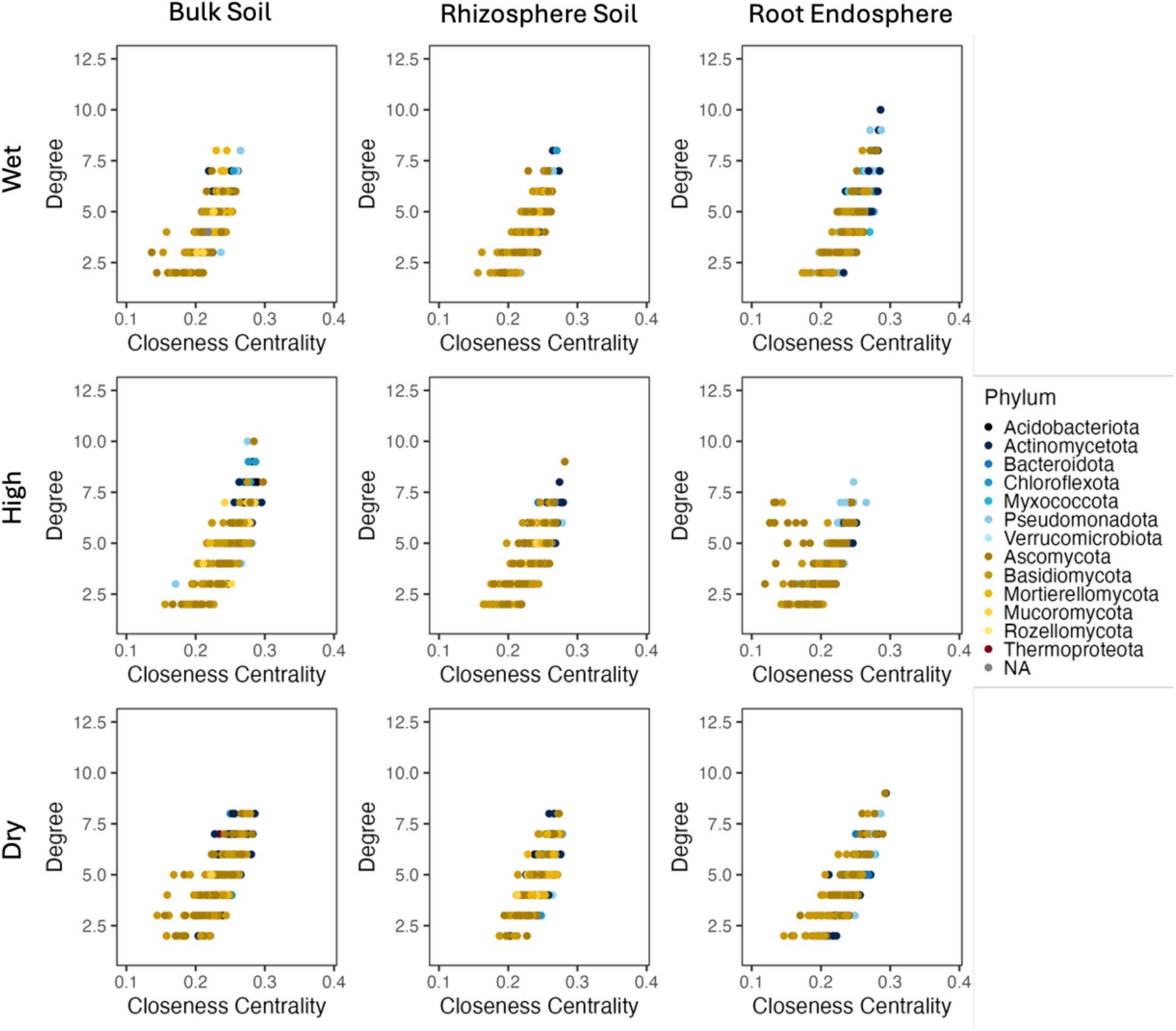
Graphs showing the degree (number of edges) vs. the closeness centrality of each node within the network. Every point represents a node within the network and is color-coded by phylum.

**S24:**
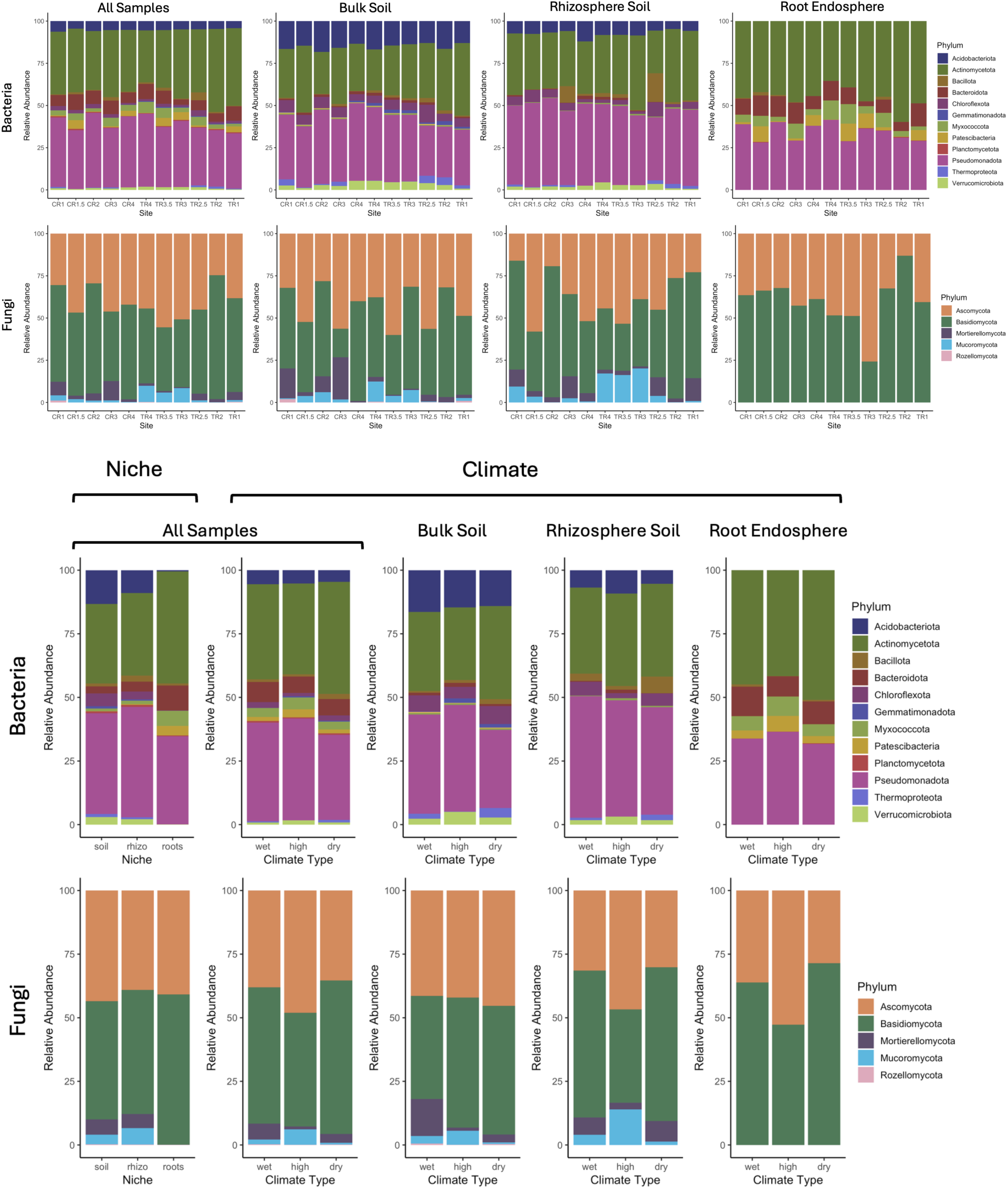
Stacked bar charts representing the relative abundance of the bacterial/archaeal (top) and fungal (bottom) phyla of all of the samples, the bulk soil samples, the rhizosphere soil samples, and the root endosphere samples (appearing left to right) across the sites, niche compartment, and climate category. Rare taxa were excluded from this analysis by trimming ASVs with fewer than 1000 reads.

**S25:**
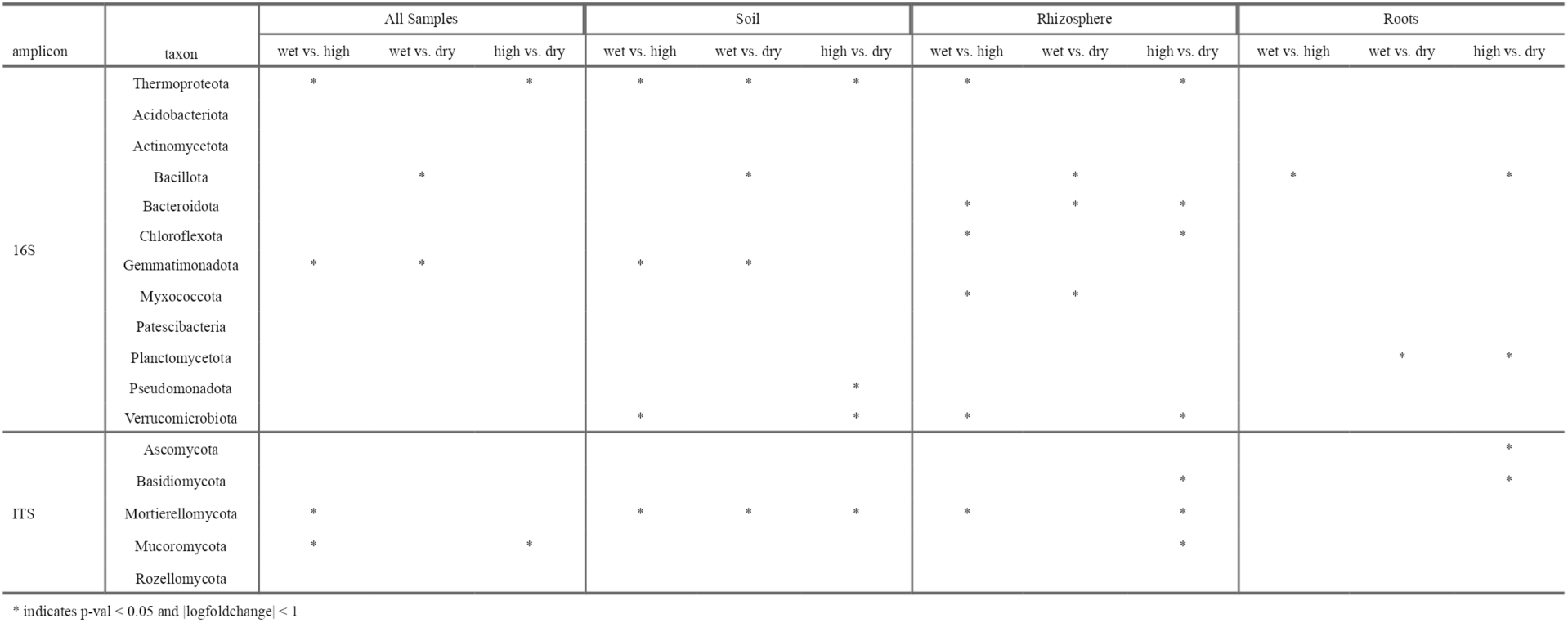
Analysis of compositions of microbiomes with bias correction (ANCOM-BC) significance tests to determine significant changes in relative abundance. Asterix indicates significant enrichment or depletion of the phylum compared to other climate classes.

**S26:** Table showing the proportion of bacterial turnover within each climate and compartment attributed to selection (heterogeneous selection and homogenous selection) and stochasticity (dispersal limitation, homogenizing dispersal, and drift and others). Selection and stochasticity columns represent the combined processes for each category. Analyses were performed in iCAMP.

| Niche | Climate | Heterogeneous Selection | Homogeneous Selection | Dispersal Limitation | Homogenizing Dispersal | Drift and Others | Selection | Stochasticity |
| --- | --- | --- | --- | --- | --- | --- | --- | --- |
| Bulk Soil | dry | 0.016 | 0.356 | 0.388 | 0.039 | 0.202 | 0.372 | 0.628 |
|  | high | 0.011 | 0.371 | 0.332 | 0.040 | 0.246 | 0.382 | 0.618 |
|  | wet | 0.008 | 0.307 | 0.386 | 0.047 | 0.253 | 0.315 | 0.685 |
| Rhizosphere Soil | dry | 0.022 | 0.343 | 0.355 | 0.041 | 0.239 | 0.365 | 0.635 |
|  | high | 0.010 | 0.369 | 0.304 | 0.035 | 0.282 | 0.379 | 0.621 |
|  | wet | 0.008 | 0.329 | 0.337 | 0.036 | 0.290 | 0.338 | 0.662 |
| Root Endosphere | dry | 0.020 | 0.413 | 0.213 | 0.013 | 0.342 | 0.433 | 0.567 |
|  | high | 0.008 | 0.326 | 0.229 | 0.016 | 0.420 | 0.334 | 0.666 |
|  | wet | 0.004 | 0.295 | 0.217 | 0.024 | 0.461 | 0.299 | 0.701 |

**S27:**
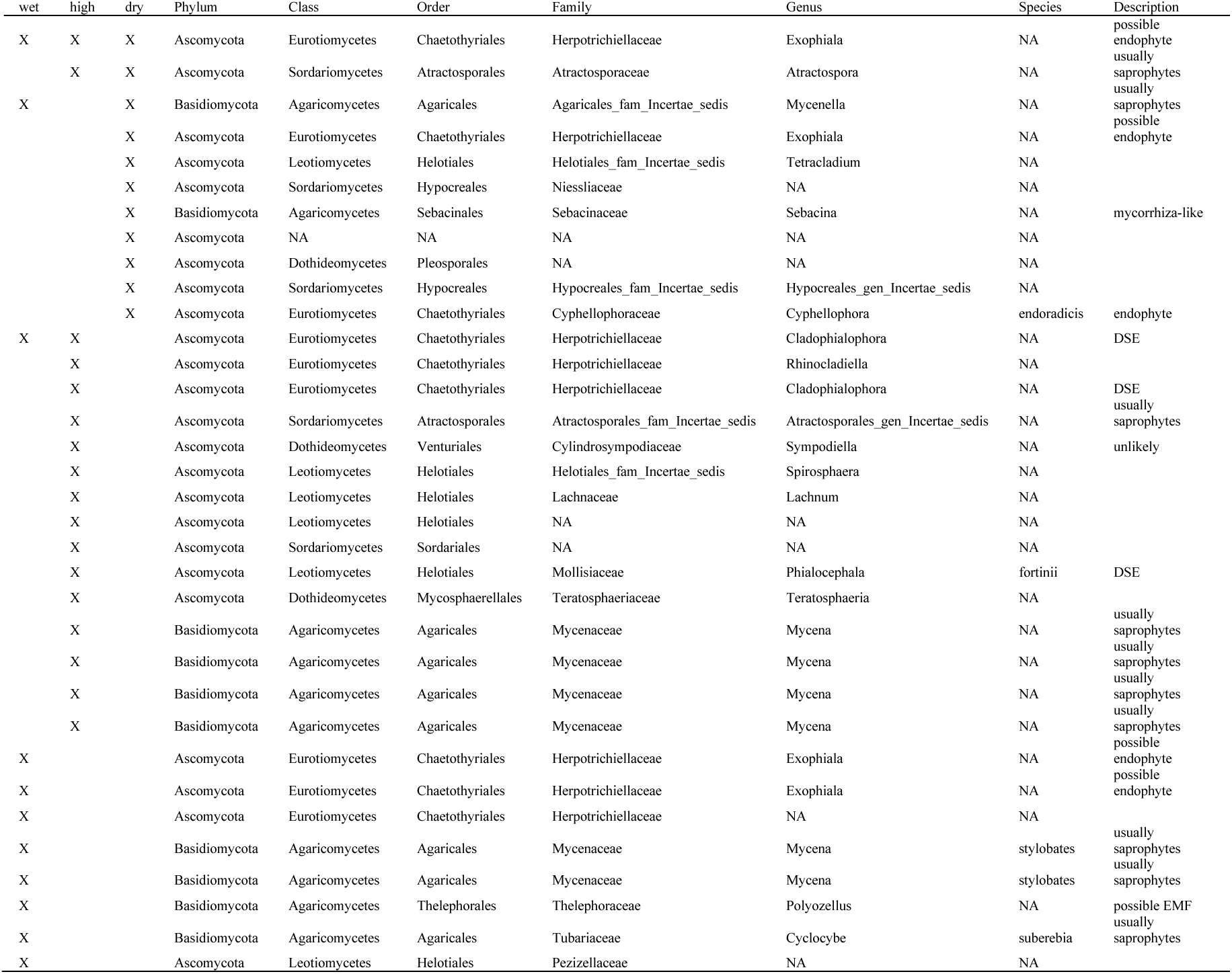
List of fungal ASVs found only in the root endosphere. Xs indicate which treatment fungi were found in. Description contains predicted function.

